# Modified base-binding EVE and DCD Domains Implicated in the Origins of Programmed Cell Death and the piRNA Pathway

**DOI:** 10.1101/2020.08.06.240630

**Authors:** Ryan T. Bell, Yuri I. Wolf, Eugene V. Koonin

## Abstract

**Background:** DNA and RNA of most cellular life forms and many viruses contain an expansive repertoire of modified bases. The modified bases play diverse biological roles that include both regulation of transcription and translation, and protection against restriction endonucleases and antibiotics. Modified bases are often recognized by dedicated protein domains. However, the elaborate networks of interactions and processes mediated by modified bases are far from being completely understood.

**Results:** We present a comprehensive census and classification of EVE domains that belong to the PUA/ASCH domain superfamily and bind various modified bases in DNA and RNA. Prokaryotes encode two classes of EVE domain proteins, slow-evolving and fast-evolving. The slow-evolving EVE domains in α-proteobacteria are embedded in a conserved operonic context that implies involvement in coupling between translation and respiration, in particular, cytochrome c biogenesis, potentially, via binding 5-methylcytosine in tRNAs. In β and γ-proteobacteria, the conserved associations implicate the EVE domains in the coordination of cell division, biofilm formation, and global transcriptional regulation by non-coding 6S small RNAs, which are potentially modified and bound by the EVE domains. Down-regulation of the EVE-encoding operons might cause dormancy or programmed cell death (PCD). In eukaryotes, the EVE-domain-containing THYN1-like proteins appear to inhibit PCD and regulate the cell cycle, likely, via binding 5-methylcytosine and its derivatives in DNA and/or RNA. Thus, the link between PCD and cytochrome c that appears to be universal in eukaryotes might have been inherited from the α-proteobacterial, proto-mitochondrial endosymbiont and, unexpectedly, could involve modified base recognition by EVE domains. In numerous prokaryotic genomes, fast-evolving EVE domains are embedded in defense contexts, including toxin-antitoxin modules and Type IV restriction systems, all of which can also induce PCD. These EVE domains likely recognize modified bases in invading DNA molecules and target them for restriction. We additionally identified EVE-like prokaryotic Development and Cell Death (DCD) domains that are also implicated in defense functions including PCD. This function was inherited by eukaryotes but, in animals, the DCD proteins apparently were displaced by the extended Tudor family, whose partnership with Piwi-related Argonautes became the centerpiece of the piRNA system.

**Conclusions:** Recognition of modified bases in DNA and RNA by EVE-like domains appears to be an important, but until now, under-appreciated, common denominator in a variety of processes including PCD, cell cycle control, antivirus immunity, stress response and germline development in animals.

## Background

Apoptosis, a poetic term derived from the Ancient Greek for petals falling off flowers or leaves falling off trees, is also referred to as programmed cell death (PCD), which conveys the biological role of the process (Kerr, Wyllie, and Currie 1972; Elmore 2007; Nagata 2018; Fuchs and Steller 2015). This phenomenon has been characterized in many types of adult metazoan tissues, as well as during embryonic development, where it is an essential counterweight to mitosis (Elmore 2007; Suzanne and Steller 2013). Defects in PCD can lead to cancer, immunosuppression and autoimmune disorders, neurodegeneration, and ischemic heart disease (Elmore 2007). Different forms of PCD with varying degrees of similarity to animal apoptosis have also been observed in diverse eukaryotes, both multicellular and unicellular (Martí;nez-Fábregas et al. 2014; Madeo et al. 2004). Moreover, evidence has accumulated that, in prokaryotes, antivirus defense systems, such as abortive infection and restriction-modification modules, as well as CRISPR-Cas, can cause PCD or dormancy (Koonin and Zhang 2017; Lewis 2000; Rice and Bayles 2003; Tanouchi et al. 2013; Makarova et al. 2012; Durand, Sym, and Michod 2016). The PCD in unicellular eukaryotes and prokaryotes is perceived as a form of altruistic suicide that prevents exhaustion of limited resources and/or abrogates the reproduction of viruses when immune mechanisms fail, promoting the survival of surrounding members of a population (Nagamalleswari et al. 2017; Fröhlich and Madeo 2000; Koonin and Krupovic 2019).

In eukaryotes, the morphological hallmarks of cells undergoing apoptosis are cytoplasmic shrinkage, chromatin condensation, fragmentation of intracellular organelles including the nucleus, plasma membrane blebbing, and ultimately, disintegration of the cell into apoptotic bodies (Elmore 2007; Kerr, Wyllie, and Currie 1972; He, Lu, and Zhou 2009; Fuchs and Steller 2015; Nagata 2018). At the molecular level, studies conducted mostly in metazoan model systems have delineated two major pathways, known as intrinsic and extrinsic. In the intrinsic pathway, the sequence of events is centered around mitochondria that integrate signals of stress or damage, and in response, release proteins from the intermembrane space into the cytosol to initiate PCD (Elmore 2007; Saikia et al. 2014; Martí;nez-Fábregas et al. 2014; Hüttemann et al. 2011). Foremost among these proteins is the heme-containing protein cytochrome c, an essential component of the respiratory electron transport chain (Kranz et al. 2009; Hüttemann et al. 2011; Ow et al. 2008; Martí;nez-Fábregas et al. 2014). Cytosolic cytochrome c binds to apoptotic protease activating factor 1 (Apaf-1), which then recruits pro-caspase-9 to assemble a multi-subunit complex, the apoptosome, starting a complex cascade of proteolytic caspase activity that results in massive protein degradation, internucleosomal DNA cleavage, and global mRNA decay (He, Lu, and Zhou 2009; Thomas et al. 2015).

The extrinsic pathway, exemplified by the interaction of cytokines of the tumor necrosis factor (TNF) family with the cognate receptors, is not initiated by mitochondrial proteins, but ultimately activates and is amplified by the intrinsic pathway (Wajant, Pfizenmaier, and Scheurich 2003; Nagata 2018; Fuchs and Steller 2015). The evolutionary precedence of the intrinsic pathway is supported by the presence of orthologs of the key apoptotic proteins in most eukaryotes as well as many bacteria (Koonin and Aravind 2002; Martí;nez-Fábregas et al. 2014; Madeo et al. 2004). In at least some unicellular eukaryotes, metacaspases, the apparent ancestors of animal caspases, have been shown to contribute to PCD (Tsiatsiani et al. 2011; Choi and Berges 2013).

Lymphocytes have been one of the primary systems employed to study PCD because PCD is required for lymphocyte development and is often disrupted in lymphomas (Rathmell and Thompson 2002). Thymocyte nuclear protein 1 (THYN1) was identified among about 300 previously uncharacterized genes that are preferentially expressed in human CD34+ hematopoietic stem/progenitor cells (Zhang et al. 2000). Shortly afterwards, a cDNA was isolated from apoptotic avian thymocytes encoding a 242 amino acid protein (Thy28) with 88% amino acid similarity to THYN1 (Compton, Thomson, and Icard 2001). Initial cloning and characterization of murine Thy28 established nuclear localization and found protein levels to be the highest in testis, with thymus, spleen, liver, and kidney also displaying substantial expression (Jiang et al. 2003). In a more recent study, nuclear THYN1 has been detected in nearly all human tissues (Thul et al. 2017).

Several studies have explored the role of THYN1/Thy28 in lymphocyte model systems where apoptosis can be induced by antibody treatment. Decreased Thy28 protein expression was observed following induction, suggesting that down-regulation of this gene is associated with apoptosis initiation (Jiang et al. 2003). Conversely, overexpression of Thy28 was correlated with inhibition of several apoptotic events, such as loss of mitochondrial membrane potential and caspase-3 activation (Toyota et al. 2012). Furthermore, these experiments established accumulation of cells in G1 phase following Thy28 overexpression, suggesting that this protein might be involved in the regulation of cell cycle progression.

THYN1/Thy28 contains a highly conserved C-terminal region now known as the EVE (named for Protein Data Bank (PDB) structural identifier 2eve) domain with readily detectable homologs in a wide variety of eukaryotes and prokaryotes (Miyaji et al. 2002). Sequence and structure analyses have shown that the EVE domain is a member of the PUA (<u>p</u>seudo<u>u</u>ridine synthase and <u>a</u>rchaeosine transglycosylase)/ASCH (<u>ASC</u>-1 <u>h</u>omology) superfamily, a widely disseminated and apparently ancient assemblage of nucleic acid-binding domains (Song et al. 2005; Yu et al. 2009; Bertonati et al. 2009; Iyer, Burroughs, and Aravind 2005; Aravind and Koonin 1999; Pérez-Arellano, Gallego, and Cervera 2007). These domains are generally associated with the translation apparatus, often fused to RNA modification enzymes, and bind RNA themselves (Pérez-Arellano, Gallego, and Cervera 2007; Iyer, Burroughs, and Aravind 2005; Aravind and Koonin 1999; Kim et al. 2017). Some ASCH domains have also been predicted to bind modified bases (Iyer et al. 2013).

In the case of EVE, THYN1/Thy28 has been characterized as a reader of 5-methylcytosine (5mC) and 5-hydroxymethylcytosine (5hmC) in DNA, as well as further oxidized 5mC derivatives 5-formylcytosine (5fC) and 5-carboxylcytosine (5caC) (Spruijt et al. 2013). Most eukaryotes encode apparent orthologs of Thy28/THYN1 in which EVE is the only recognized domain, although fusions with AT-hook and other domains in fungi have been described, further supporting the role of EVE as a DNA-binding domain in these proteins (Iyer et al. 2013). The PUA-like SRA (<u>S</u>ET and <u>R</u>ING-<u>a</u>ssociated) domain also binds 5mC and 5hmC DNA (Hashimoto et al. 2008; Spruijt et al. 2013), however, the initial description of the EVE domain identified another PUA-like domain, YTH (<u>YT</u>521-B <u>h</u>omology), as the most similar to EVE of the PUA-like structures in the PDB (Bertonati et al. 2009). YTH also binds modified bases, recognizing N^6^-methyladenosine (m^6^A) in RNA in eukaryotic proteins and m^6^A DNA in an archaeal protein (Patil, Pickering, and Jaffrey 2018; Hosford, Bui, and Chappie 2020). The conserved core of the superfamily consists of a 5-stranded β-barrel fold (green in Figure 1), often with an alpha helix between strands 1 and 2, a structural element that is present in EVE domains, which also contain an additional sixth strand in the β-barrel (Bertonati et al. 2009).

**Figure 1:**
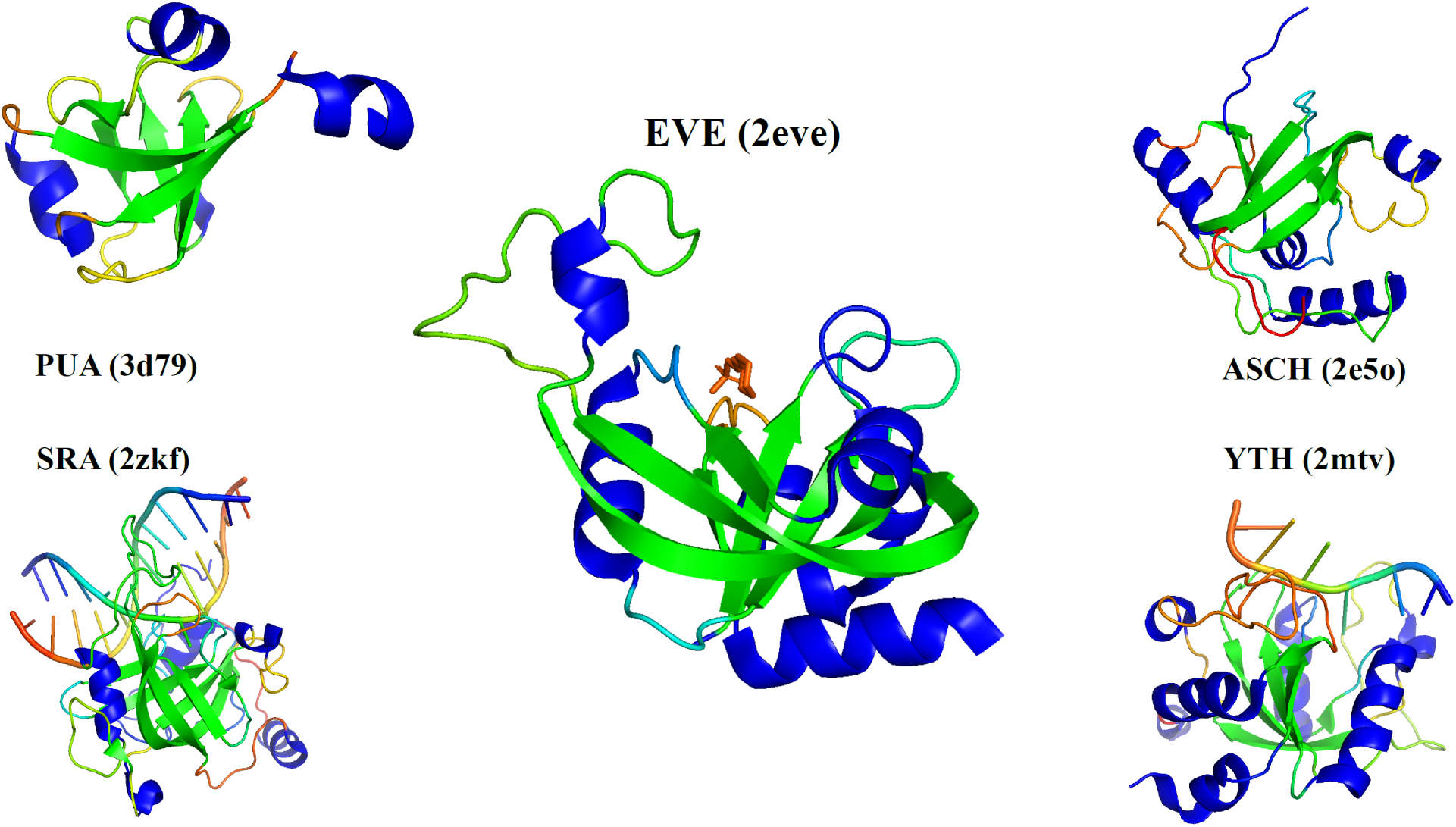
Structures of EVE and other PUA/ASCH superfamily members. Structures were downloaded from PDB (identifiers in parentheses) and drawn using the PyMOL program (Schrodinger 2015). β-strands are colored green, α-helices are colored blue, while loops and ligands are rainbow colored.

Here, we report a comprehensive bioinformatic analysis of the broad phyletic distribution of EVE-like domains, with an emphasis on the radiation among Proteobacteria, intriguing associations with base modification-dependent restriction and toxin-antitoxin systems, and the identification of the Development and Cell Death (DCD) domain as a member of the EVE-like superfamily. We apply the ‘guilt by association’ approach (Aravind 2000; Vey 2013; Rogozin et al. 2004; Doerks, von Mering, and Bork 2004; Galperin and Koonin 2000) to make functional inferences from an extensive comparative analysis of the expanded collection of bacterial and archaeal genomes.

## Results

### A census of EVE proteins

Our search for EVE proteins using PSI-BLAST and HHpred seeded with profiles derived from multiple alignments of the amino acid sequences of known EVE domains (see Methods for details) showed that the EVE domain is most prevalent among Proteobacteria, which harbor the majority of all prokaryotic EVE proteins detected (Supplementary Figure 1) and a plurality of all EVE proteins. CLANS analysis (Frickey and Lupas 2004) of EVE domains extracted from all EVE proteins in the dataset revealed a diverse cloud of sequences, with four well-defined clusters (Figure 2). The largest cluster (blue in Figure 2) consists, mostly, of sequences from β and γ-proteobacteria, as well as those from the metazoa and fungi. The second largest cluster (red) includes mostly sequences from α-proteobacteria and Bacteroidetes, as well as the majority of plant sequences. Two smaller, almost completely prokaryotic clusters were also identified. The first (green) represents a collection of sequences largely from Proteobacteria, Actinobacteria, and Bacteroidetes. These EVE domains are usually encoded in operonic contexts which imply a role in ligand-activated transcriptional regulation. The second (purple) is mostly made up of sequences from γ-proteobacteria, Firmicutes, and Bacteroidetes and is unique in that the EVE domains in this group are almost always fused to a GNAT (<u>G</u>CN5-related <u>N</u>-<u>a</u>cetyltransferase) domain.

**Figure 2:**
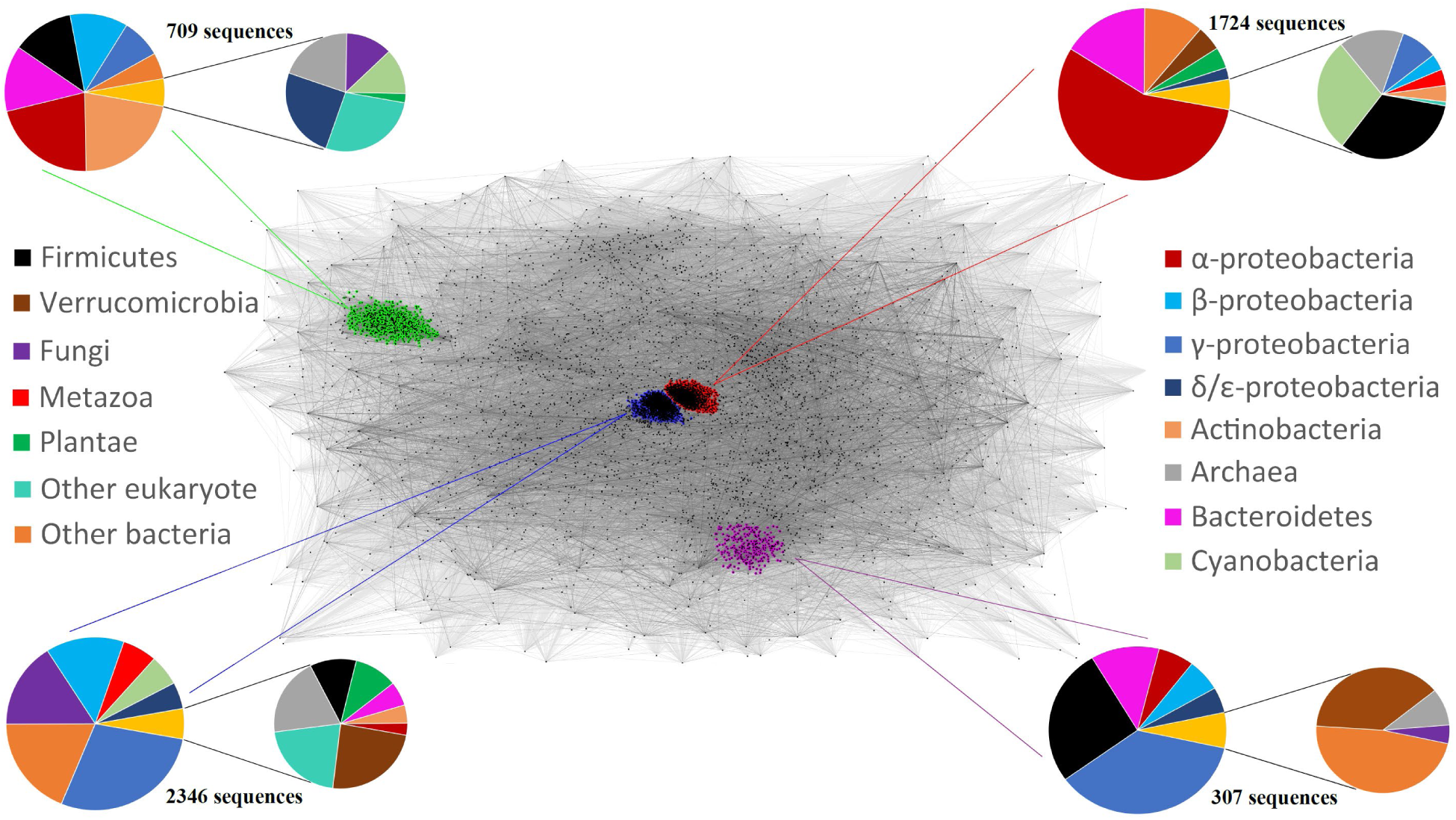
CLANS analysis of EVE domains. A 2D projection of CLANS clustering analysis of 8,403 representative EVE domain sequences. Each sequence is depicted by a dot and sequence similarity detected by BLAST is indicated by a line, colored in shades of gray according to the BLAST p-value. Four primary clusters were observed, marked with colors which match colored lines extending toward the pie chart which corresponds to the cluster. The protein sequences used for the CLANS analysis are available as Supplementary Dataset 1.

We chose to focus our initial analysis on the two large clusters, which consist, mostly, of proteobacterial EVE domains. α-proteobacteria were the most abundant class in the data, from which the majority of sequences in the second largest cluster (red in Figure 2) derive.

### EVE in α-proteobacteria

The EVE proteins of this class (Figure 3) are frequently located in a putative operon with the tRNA N^6^-adenosine threonylcarbamoyltransferase TsaD, glycerol-3-phosphate dehydrogenase GpsA, and YciI, a small ferredoxin-fold protein homologous to muconolactone isomerases (Willis et al. 2005). The sequences of the EVE domains in this group are readily recognizable (RPS-BLAST E-values of ∼1e-42 or better with the pfam01878 query) and form a tight, well-conserved collection with within-group divergence comprising only 35% of the overall divergence between EVE domains (see Methods for details).

**Figure 3:**
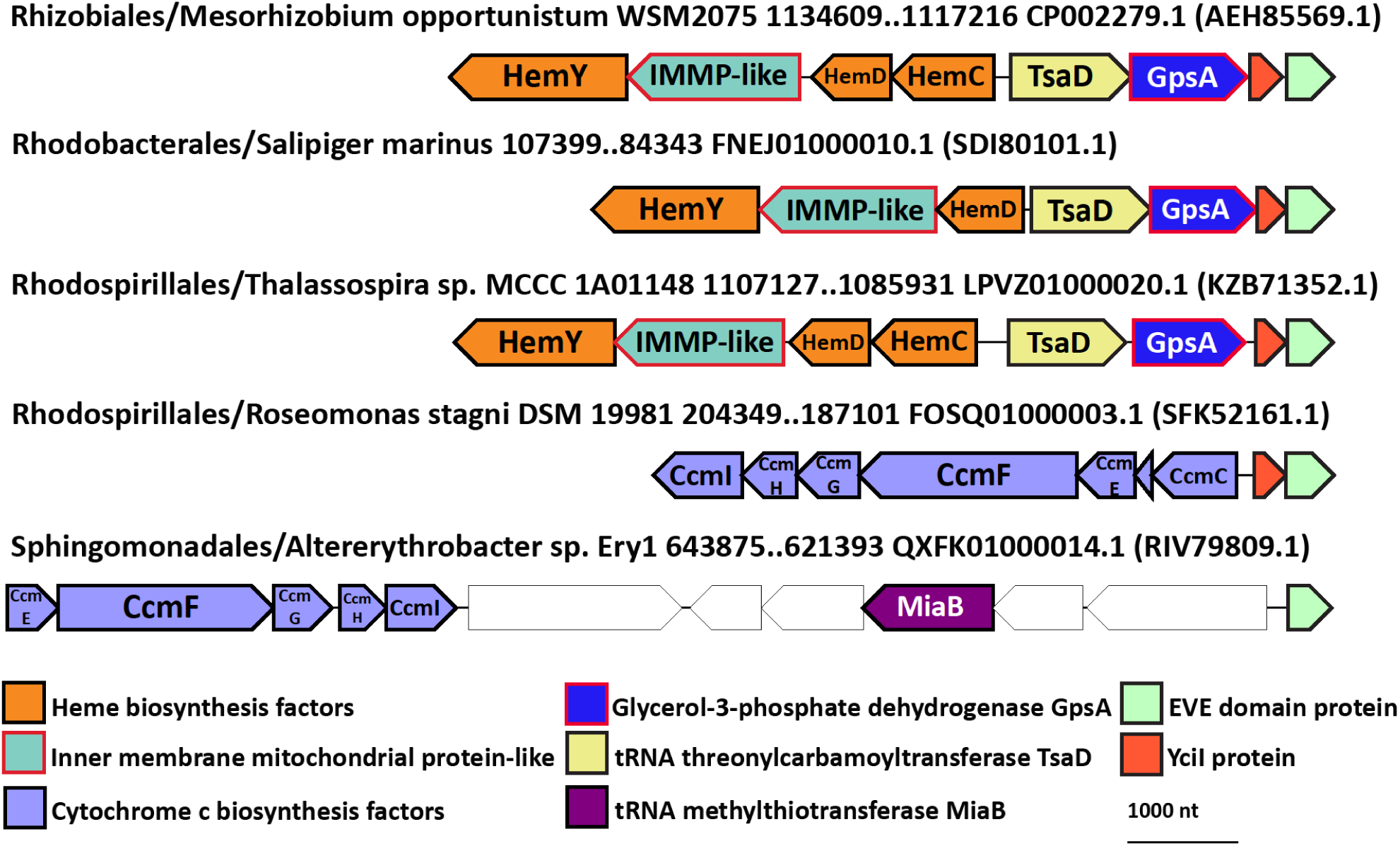
Conserved genomic context of EVE proteins in α-proteobacteria. Representative EVE protein neighborhoods from α-proteobacteria. Genes are shown as arrows from 5’ to 3’. The order of α-proteobacteria, species, and genomic coordinates for each neighborhood are indicated, as are the GenBank genome accessions and, in parentheses, the GenBank accessions for each EVE protein.

This highly conserved directional unit (TsaD->GpsA->YciI->EVE) is itself strongly associated with another predicted operon which encodes 3 enzymes of heme biosynthesis, namely, porphobilinogen deaminase (HemC), uroporphyrinogen-III synthase (HemD), and coproporphyrinogen oxidase (HemY/HemG), as well as a diverged homolog of HemX, a putative uroporphyrinogen-III C-methyltransferase that is also homologous to IMMP (inner <u>m</u>embrane <u>m</u>itochondrial <u>p</u>rotein, also known as mitofilin) (Huynen et al. 2016). In Rhodobacteraceae, HemC is missing from this generally well conserved gene order. Head to head orientation of these putative operons suggests that the promoter regions might overlap, allowing for co-regulation.

The association between the EVE domain and cytochrome c biosynthesis via regulation of heme production in α-proteobacteria is further emphasized by the presence of a cytochrome c biosynthetic cluster (CcmC through CcmI) adjacent to the EVE domain that is conserved in both the Acetobacteraceal branch of the Rhodospirillales and the Sphingomonadales (Figure 3) (Kranz et al. 2009). In Sphingomonadales, a likely operon including the tRNA modifying enzyme MiaB, which adds a methylthio group to *N*^6^-isopentenyladenosine at position 37 in many tRNAs decoding UNN (the same position modified by TsaD), often occurs between the EVE domain and the cytochrome c biosynthetic operon (Pierrel et al. 2004).

A contextual information network graph generated from the pairwise domain associations in prokaryotic EVE protein genomic neighborhoods showed that in α, β, and γ-proteobacteria, respectively, the EVE proteins are associated with highly conserved, but largely non-overlapping gene complements (Figure 4).

**Figure 4:**
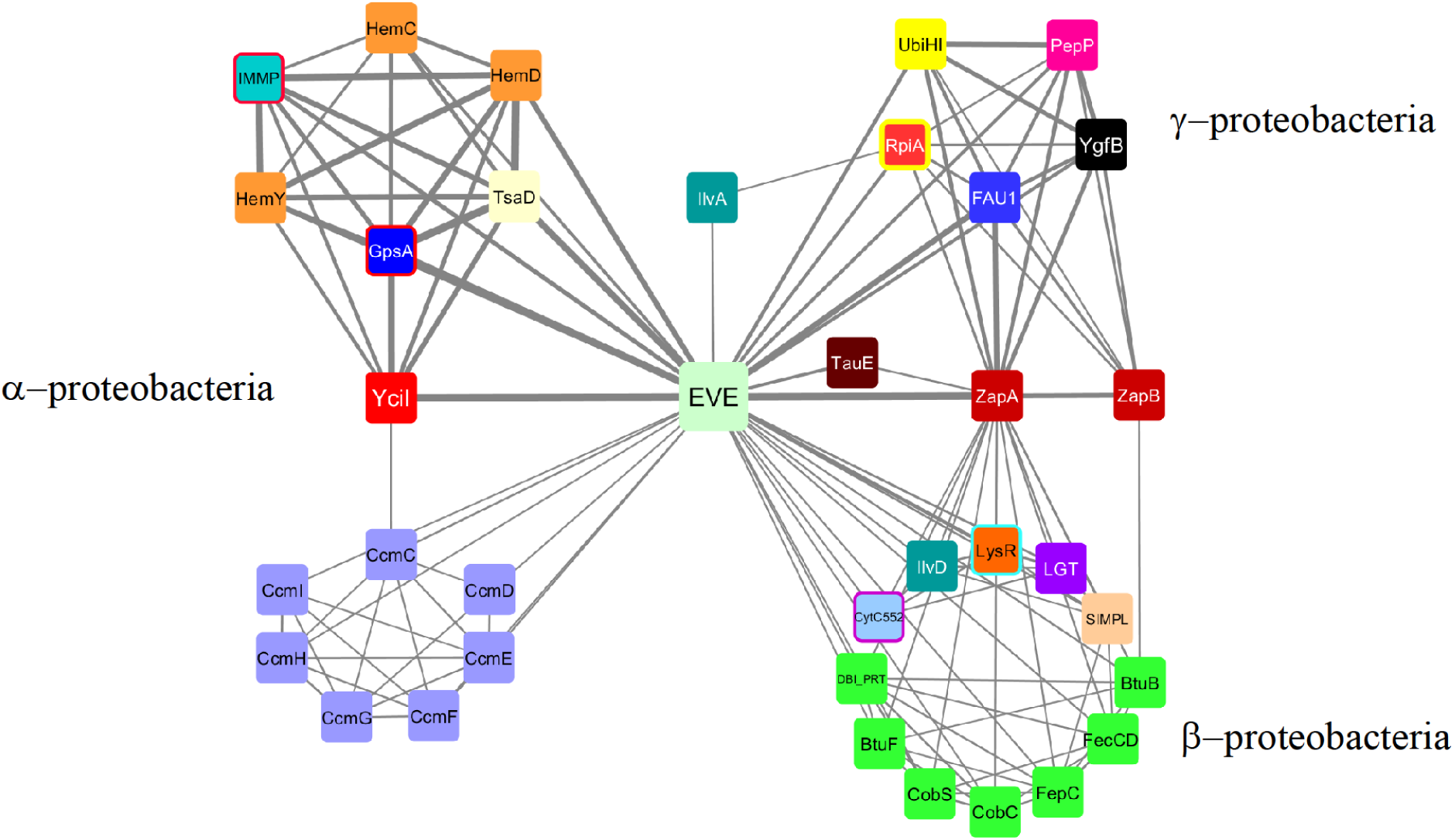
The most prevalent contextual associations of EVE proteins in the genomes of Proteobacteria. A contextual information network graph of domains detected among genes in the genomic neighborhoods encoding EVE proteins in Proteobacteria. The nodes are clustered by mutual connections, which generally correlate with the distribution of species in which they occur. The thickness of the edges reflects the strength of the association. Nodes representing domains that participate in the same pathway are drawn with the same color. Domain coloring and abbreviations are explained in the text and Figures 3 and 5. The graph was calculated from the top 400 pairwise associations between domains in EVE protein neighborhoods from 13,388 genomes with unique domain compositions. Singleton associations with EVE and minor networks were removed for clarity.

### EVE in β and γ-proteobacteria

A prominent exception to the general lack of overlap between the contextual information networks among Proteobacteria is the conservation between β and γ-proteobacteria of an apparent operonic linkage of EVE proteins and the cell division proteins ZapA and ZapB (Figure 4). The sequences of the EVE domains in these proteins are also highly recognizable, slightly more so, in fact, than those in α-proteobacteria (RPS-BLAST E-values of ∼5e-55 or better). They likewise form a tight, well-conserved group, with their within-group divergence accounting for only 32% of the overall divergence between EVE domains. The protein-coding gene array ZapB->ZapA->EVE also contains, between ZapA and EVE, a non-coding 6S RNA (*ssrS*) gene. Our analysis of these neighborhoods suggests that the *ssrS* gene is (nearly) always present, based on the positions of the protein-coding genes, leaving a gap sufficient to accommodate the 6S RNA, but are not consistently annotated, conceivably, due to sequence divergence. For this reason, *ssrS* was not included in our calculations that produced the contextual information network graph (Figure 4).

In many species of γ-proteobacteria and some β-proteobacteria, the enzyme FAU1/MFTHFS, also known as YgfA, a putative 5-formyltetrahydrofolate cyclo-ligase, is encoded between ZapB->ZapA->SsrS and the EVE protein (Figure 5). In γ-proteobacteria, another directional gene array is frequently found adjacent to this predicted operon in a head to head orientation, with the potential for the promoter regions to overlap. It encodes an uncharacterized conserved protein (YgfB), an Xaa-Pro aminopeptidase (PepP), and a homolog of 2-octaprenyl-6-methoxyphenol 4-hydroxylase (UbiH), an FAD-dependent oxidoreductase, as well as a homolog of 2-octaprenylphenol 6-hydroxylase (UbiI), both of which are involved in ubiquinone biosynthesis (Figure 5) (Meganathan 2001). Many of the γ-proteobacterial neighborhoods additionally include genes encoding homologs of ribose-5 phosphate isomerase (RpiA) and L-threonine dehydratase (IlvA).

**Figure 5:**
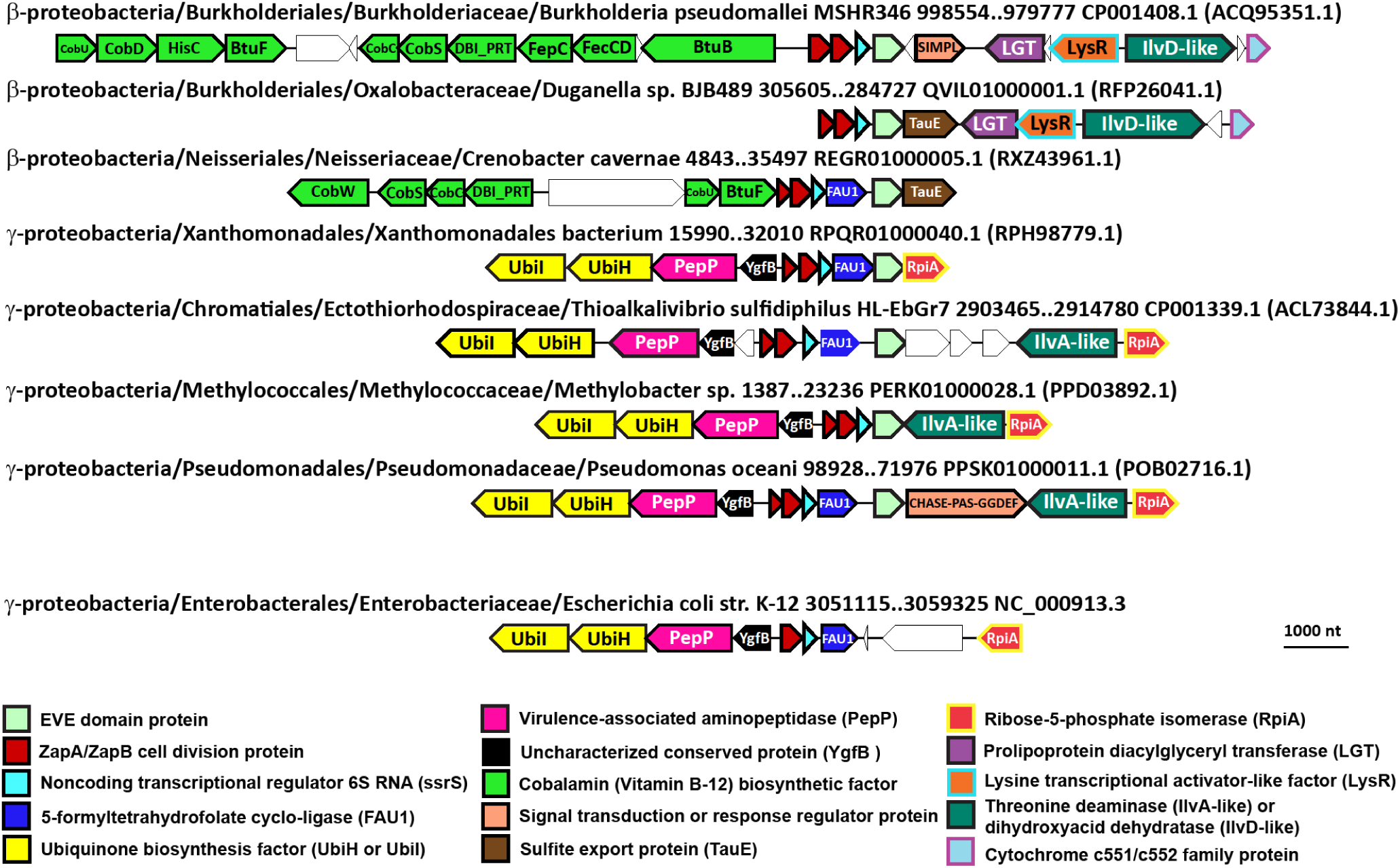
Conserved genomic context of EVE proteins in β and γ-proteobacteria. Representative EVE protein neighborhoods from β and γ-proteobacteria. Genes are shown as arrows from 5’ to 3’. The taxonomic lineage and genomic coordinates for each neighborhood are indicated, as are the GenBank genome accessions and, in parentheses, the protein accessions for each EVE protein. A homologous region from the genome of *E. coli* K-12 that lacks the EVE domain protein is shown for reference at the bottom of the figure.

In β-proteobacteria, the gene coding for the EVE protein is often followed by a gene encoding the ortholog of the TauE sulfite export protein (Figure 5). In Burkholderiaceae and Neisseriales, a cobalamin (Vitamin B-12) biosynthetic cluster is often found immediately adjacent to the ZapAB->SsrS->(FAU1)->EVE unit (Figure 5). In Burkholderiales, a conserved region encoding a cytochrome c551/c552 family protein, dihydroxy-acid dehydratase (IlvD), a putative transcriptional regulator related to LysR, and prolipoprotein diacylglyceryl transferase (LGT) is adjacent to the ZapAB->SsrS->EVE putative operon.

The recently updated phylogeny of the β and γ-proteobacteria (Williams et al. 2010) allows some inferences to be made concerning the evolutionary history of the predicted functional systems containing the EVE domain. The taxonomic distribution of the ZapAB->SsrS->(FAU1)->EVE unit covers β-proteobacteria, several early branching members of γ-proteobacteria (Xanthomonadales, Chromatiales, Methylococcales, etc.), and the clade primarily consisting of Pseudomonadales and Oceanospirillales. This broad taxonomic representation implies that the unit was present in the common ancestor of β and γ-proteobacteria. The VAAP clade (Vibrionales, Alteromonadales, Aeromonadales, and Pasteurellales) have lost this association, and each order, with the exception of Aeromonadales, possesses distinct conserved regions neighboring encoded EVE proteins (Figure 12, Supplementary Figures 2-4). In *E. coli* K-12, both of the typical EVE-associated γ-proteobacterial operons and their orientations are conserved, but the ZapB and EVE domain proteins have been lost (Figure 5).

In agreement with the CLANS results, in the phylogenetic tree of the EVE domains, the EVE proteins from most of the higher plants branch from within the α-proteobacterial clade, whereas EVE proteins from the metazoa, fungi, and some plants are more similar to γ-proteobacterial domains, but lie outside of the γ-proteobacterial variation (Supplementary Figure 5). Due to the small size of the EVE domain, phylogenetic analysis cannot confidently identify the prokaryotic ancestry of these domains in eukaryotes, although Proteobacteria are the most likely contributors, with possible multiple acquisitions.

### EVE domains in putative ligand-activated antibiotic resistance and other ligand-activated responses

The largest of the almost exclusively prokaryotic clusters from our CLANS analysis (green in Figure 2) was populated predominantly by domains encoded in the operonic context of a transcription factor and a small molecule ligand-binding domain (Figure 6). The most frequent putative operons encoded an EVE domain with either a MarR (<u>m</u>ultiple <u>a</u>ntibiotic resistance) family transcription factor or a YafY family transcription factor. YafY-like factors are a fusion of a putative DNA-binding HTH domain and a WYL domain, a ligand-binding regulator of prokaryotic defense systems (Makarova et al. 2014; Yan et al. 2018; Müller et al. 2019). The MarR-EVE and YafY-EVE pairs are further associated, most frequently, with a ligand-binding domain of the SPRBCC (<u>S</u>TART/<u>R</u>HO_alpha_C/<u>P</u>ITP/<u>B</u>et_v1/<u>C</u>oxG/<u>C</u>alC) or EhpR (phenazine antibiotic resistance) families. EhpR family proteins contain a vicinal oxygen chelate (VOC) domain, and other VOC domain homologs are also frequently encoded in the neighborhoods of this class of EVE proteins, often replacing SPRBCC domains in association with MarR-EVE pairs (Figure 6). The sequences of this group of EVE domains formed a distinct clade in our phylogenetic analysis (Supplementary Figure 5). The regions surrounding these apparent 3-component systems are highly diverse. They include putative defense functions in *Nocardia* and related genera, where multiple paralogs of UvrD-like helicase domains fused to Cas4-like PD(D/E)XK phosphodiesterases (Hudaiberdiev et al. 2017) are present (Supplementary Figure 6). Conversely, in *Azospirillum*, the neighborhoods include translation factor genes and cytochrome c biosynthesis operons, a context that is, surprisingly, closely similar to the distinct classes of EVE proteins in the two largest clusters in our CLANS analysis (Supplementary Figure 7).

**Figure 6:**
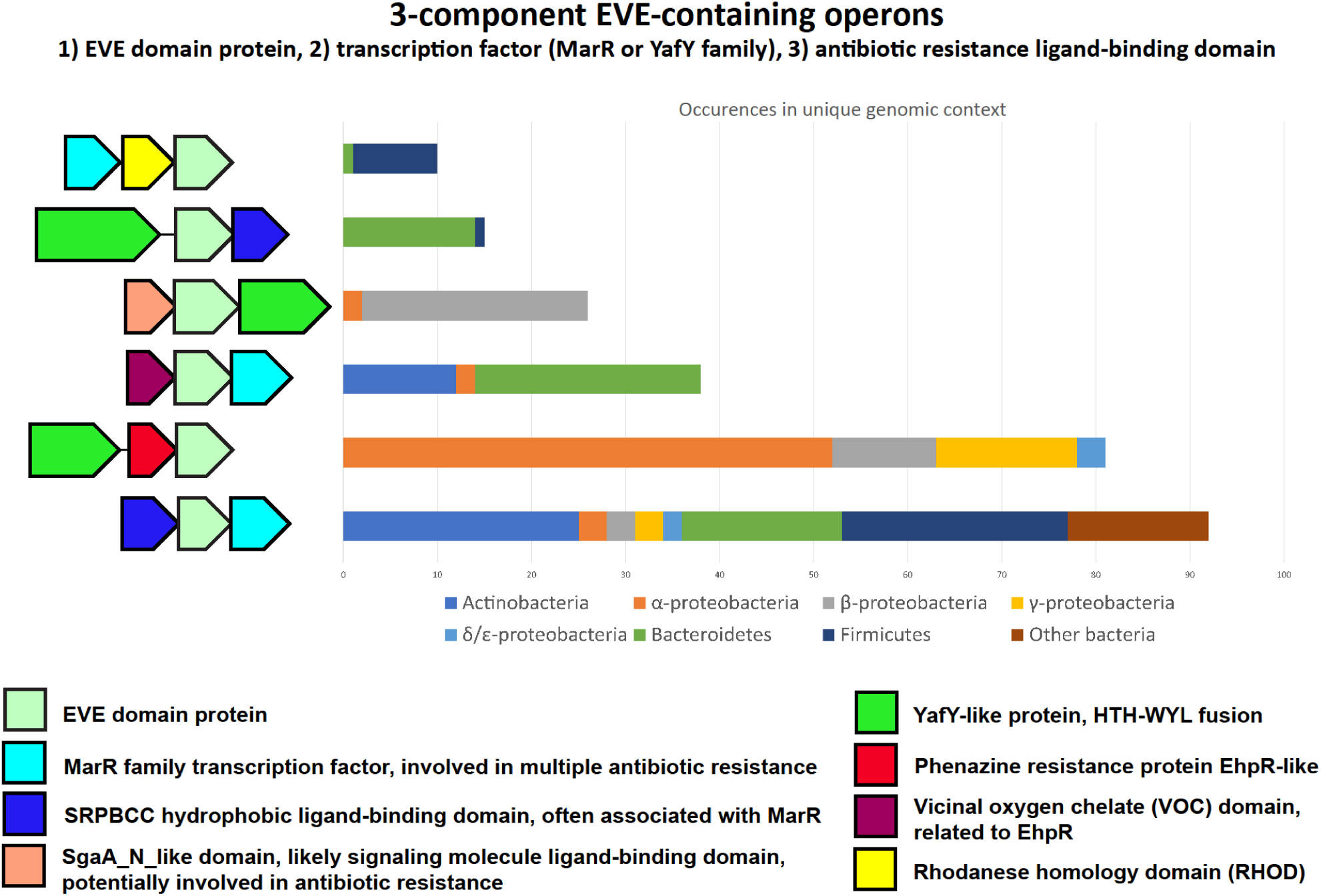
A distinct class of EVE proteins frequently encoded in the context of ligand-binding antibiotic resistance and defense regulators as well as transcription factors. Representative operons encoding EVE proteins found in the green CLANS cluster from Figure 1. Genes are shown as arrows from 5’ to 3’. The order of the genes may vary within a given group.

**Figure 7:**
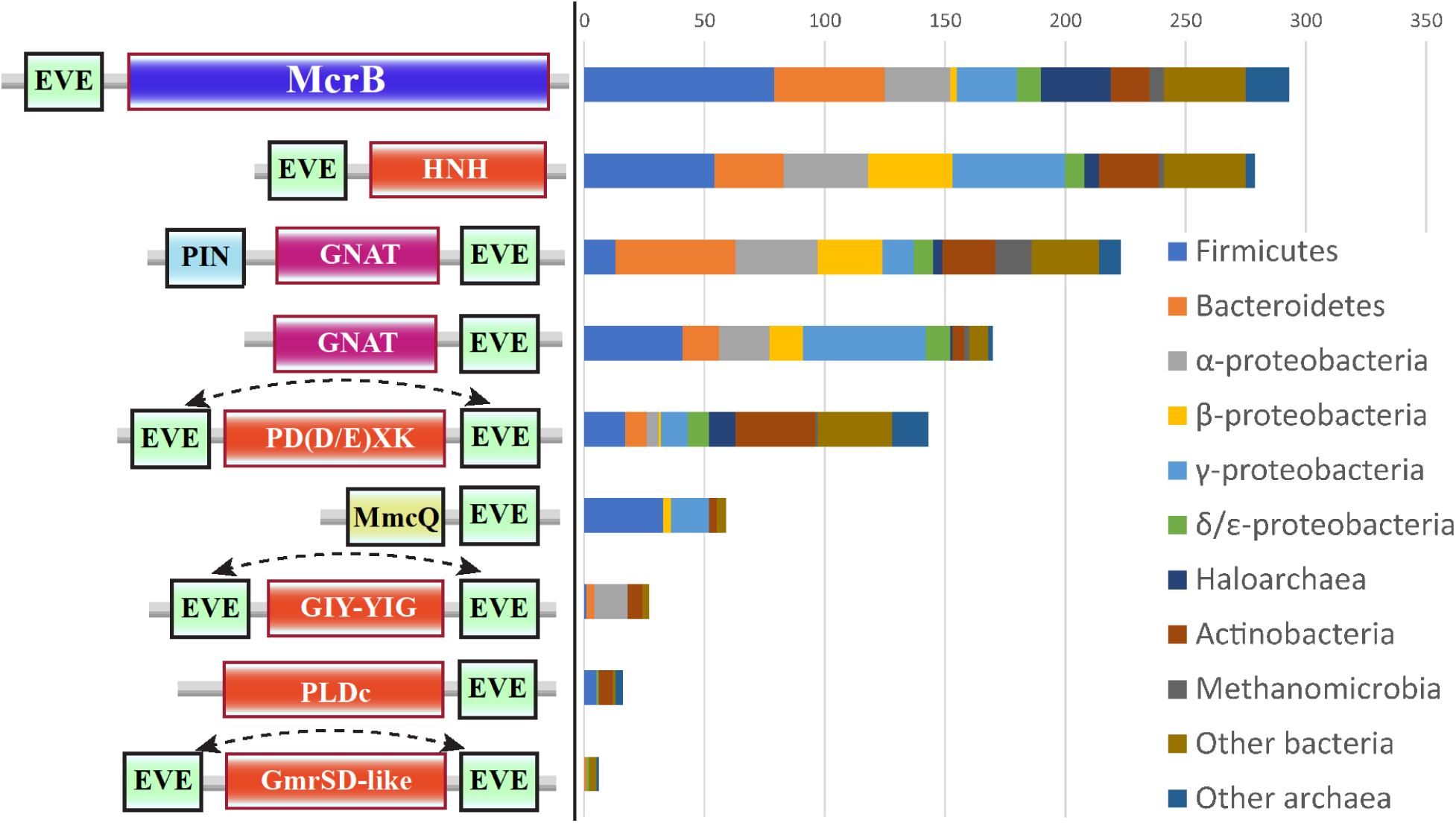
Phyletic distributions of classes of EVE fusion proteins with predicted roles in prokaryotic modification-dependent restriction and toxin-antitoxin systems. The most common classes of EVE fusion proteins in prokaryotes, with one representative chosen per genus. The representatives contain the core elements depicted (not to scale), with additional domains often present, especially in EVE-McrB proteins. The dotted lines with an arrow at each end indicate that the EVE domain can occur in either position, but not both.

### EVE as a specificity domain in modification-dependent restriction systems

The EVE proteins in Proteobacteria and eukaryotes found in the two largest clusters we observed with CLANS analysis show high levels of sequence conservation. By contrast, many genome defense systems encompass EVE domains with more pronounced sequence diversity. These domains range from highly significant matches to hits with weaker similarity, and many could be detected only with sensitive methods such as HHpred. A substantial variety of putative modification-dependent (Type IV) restriction endonucleases (REs) with core architectures of EVE-PD(D/E)XK phosphodiesterase and EVE-HNH endonuclease were identified in our searches (Figure 7). Furthermore, we identified numerous proteins containing fusions of the EVE domain with nucleases of the Phospholipase D (PLDc) or GIY-YIG superfamilies (Figure 7). Rare fusions to homologs of the glucosylated 5hmC-dependent RE GmrSD were detected as well.

The EVE domain is also frequently incorporated into homologs of the GTP-dependent DNA translocase McrB. In *E. coli* K-12, McrB, in concert with McrC, a PD(D/E)XK-type nuclease that interacts with McrB hexamers via its N-terminal domain, restricts N4-methylcytosine (4mC)/5mC/5hmC-containing DNA; in this strain, EVE is replaced with a DUF3578 family domain as the specificity module (Dila et al. 1990; Sukackaite et al. 2012; Hosford, Bui, and Chappie 2020; Nirwan et al. 2019). Overall, the EVE-McrB combination is the most common domain architecture among the EVE-containing proteins in defense systems, represented in nearly 300 bacterial and archaeal genera, and is particularly abundant among Firmicutes and Bacteroidetes (Figure 7).

In diverse archaea, a recurrent partnership was observed between standalone EVE domain proteins and a predicted, uncharacterized restriction system that encodes a SWI2/SNF2 helicase fused to a nuclease (PD(D/E)XK or PLDc family). This gene is expressed in an operon that also encodes a methyltransferase of COG1743 and an uncharacterized DUF499-containing protein (Supplementary Figure 8). Our analysis showed that DUF499 is homologous to CDC6/ORC1 ATPases, which are involved in the recognition of the origin of DNA replication in archaea and eukaryotes (Akita et al. 2010; Makarova and Koonin 2013).

**Figure 8:**
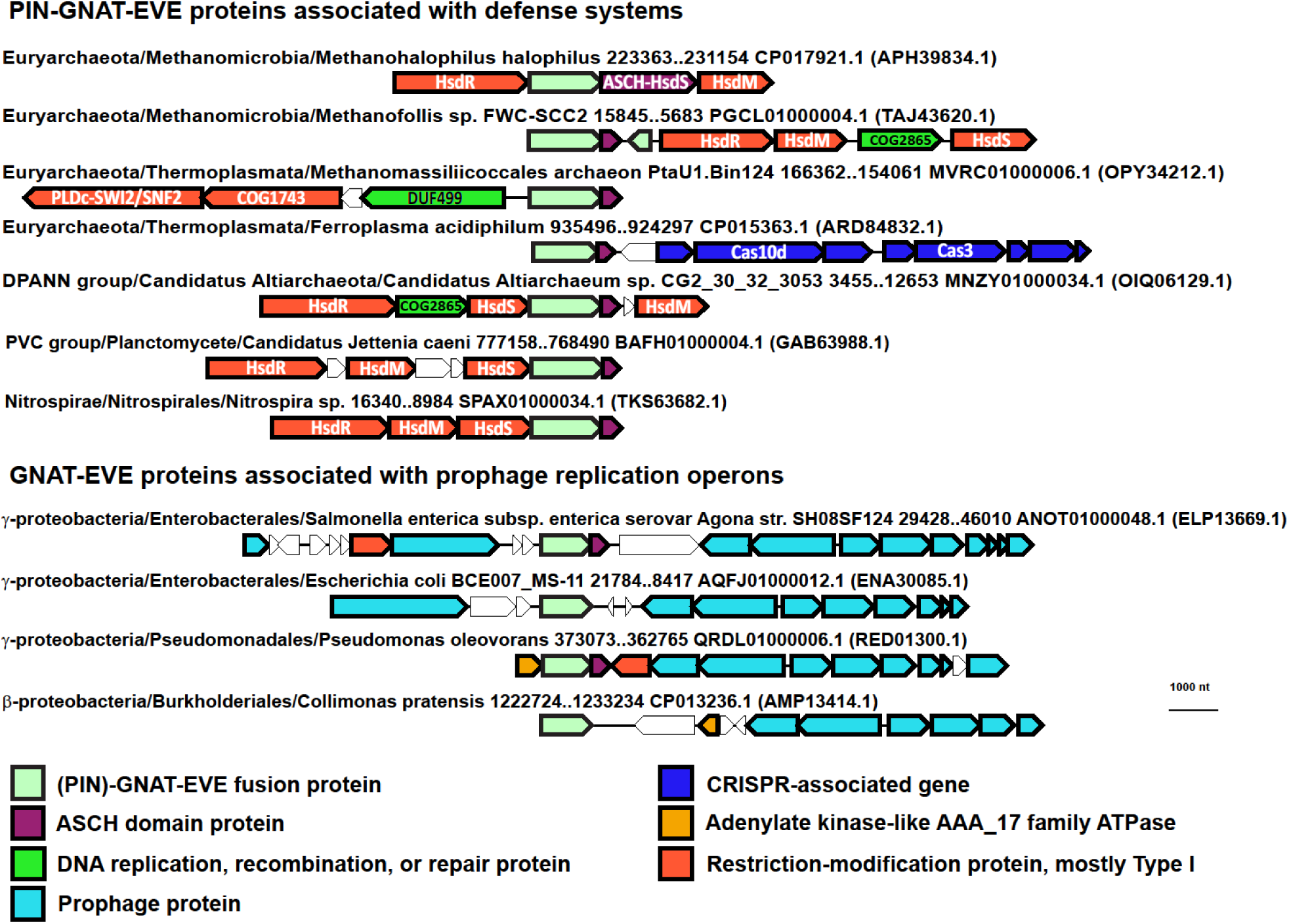
Distinct genomic associations between PIN-GNAT-EVE and GNAT-EVE proteins. Representative (PIN)-GNAT-EVE protein neighborhoods from archaea and bacteria. Genes are shown as arrows from 5’ to 3’. The taxonomic lineage and genomic coordinates for each neighborhood are indicated, as are the GenBank genome accessions and, in parentheses, the GenBank accessions for each EVE protein. COG2865 is labeled as a DNA replication, recombination, or repair protein due to similarity to RecG DNA helicase detected by HHpred (99.7% probability, E-value: 1.6e-15).

### EVE domains in toxin-antitoxin systems

A major class of EVE proteins which formed a distinct cluster in our CLANS analysis (purple in Figure 2), is a fusion of EVE to the C-terminus of a GNAT-like acetyltransferase, often with a PIN RNase domain at the N-terminus (Figure 7). GNAT and PIN domains both frequently function as toxins (Makarova, Wolf, and Koonin 2009; Matelska, Steczkiewicz, and Ginalski 2017; Yeo 2018). This variety of EVE proteins has been described previously in some detail by Iyer et al., who proposed that these proteins acetylate a DNA base, although the frequent presence of a PIN domain suggests that these systems employ RNA as a target or guide (Iyer et al. 2013). As also addressed in that study, almost all (PIN)-GNAT-EVE operons encode a protein containing a second PUA-like domain, ASCH, and often, also, an AAA+ ATPase of the AAA_17 family. In some cases, mostly in α-proteobacteria, the ASCH domain is fused to a helix-turn-helix (HTH) DNA-binding domain of the xenobiotic response element (XRE) family.

The distributions of the PIN-GNAT-EVE and the GNAT-EVE fusion proteins among prokaryotes are notably different (Figure 8). The PIN-GNAT-EVE proteins are frequently found in bacterial and archaeal genomes in a close association with Type I restriction-modification (RM) systems (HsdR/M/S operons). A consistent proximity between PIN-GNAT-EVE proteins and other types of defense systems, such as CRISPR-Cas and the COG1743->DUF499->SWI2/SNF2 helicase-nuclease operon described above, was also observed (Figure 8). By contrast, GNAT-EVE proteins are not associated with Type I RM systems but are commonly located within prophages in β and γ-proteobacteria (Figure 8).

In addition to the profusion of putative TA systems containing EVE domains, EVE is also regularly found as a standalone protein closely associated with Type I RM systems (Figure 10). These systems often also contain an ASCH domain and are mostly found in archaea. In effect, RM systems exhibit toxin-antitoxin functionality, with the restriction endonuclease playing the role of toxin, whereas the methyltransferase is its antitoxin (Mruk and Kobayashi 2014; Vasu and Nagaraja 2013; Makarova, Wolf, and Koonin 2009; Naito, Kusano, and Kobayashi 1995; Fozo et al. 2010). Accordingly, the EVE domains are likely to play similar roles in these systems, namely, targeting the toxins (including restriction endonucleases) to modified nucleic acids.

**Figure 9:**
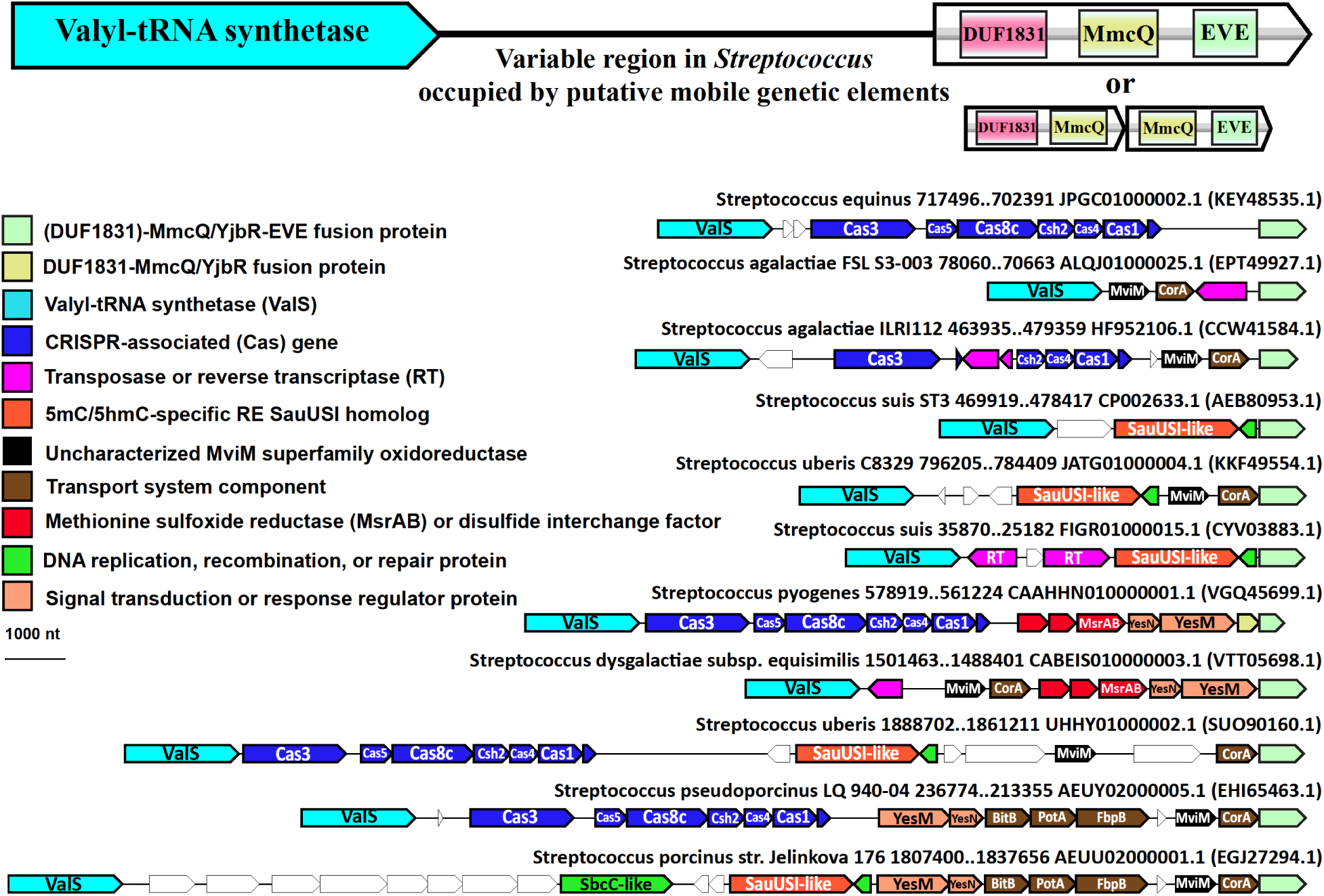
MmcQ/YjbR-EVE fusion proteins in *Streptococcus*. Representative (DUF1831)-MmcQ/YjbR-EVE protein neighborhoods from *Streptococcus*. Genes are shown as arrows from 5’ to 3’. The genomic coordinates for each neighborhood are indicated, as are the GenBank genome accessions and, in parentheses, the GenBank accessions for each EVE protein. The small ‘DNA repair’ proteins adjacent to the SauUSI homologs are homologs of MutT pyrophosphohydrolase.

**Figure 10:**
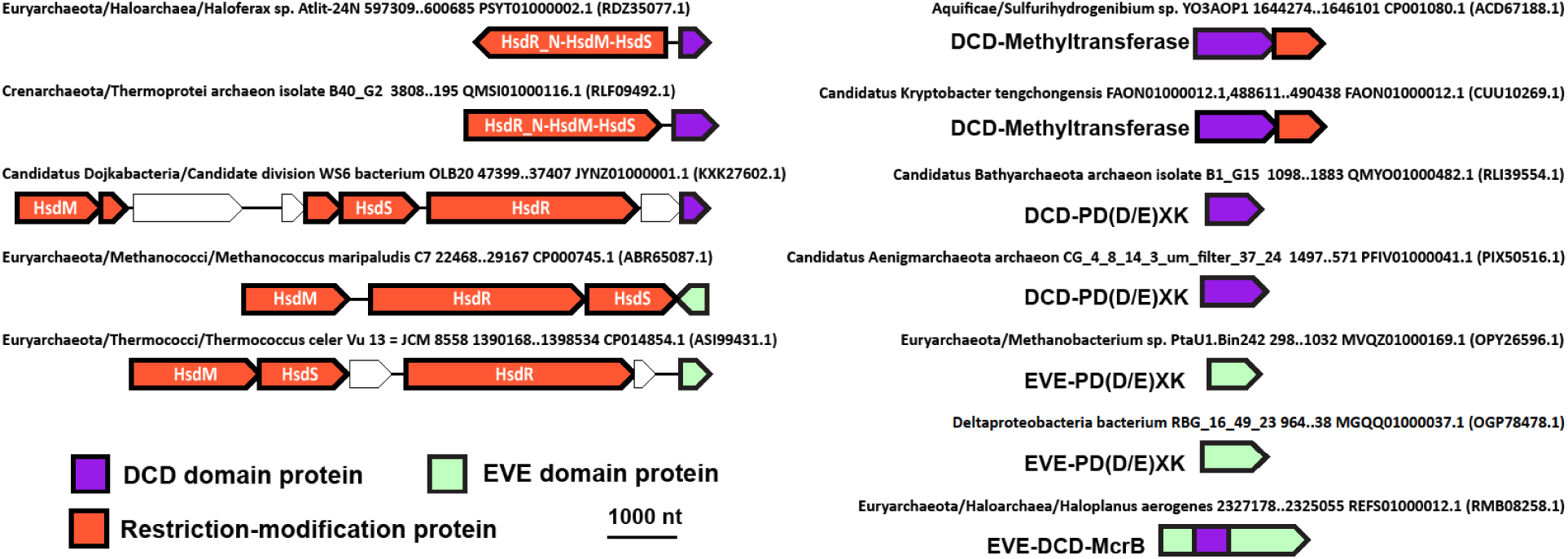
DCD is an EVE-like domain. Representative DCD and EVE protein neighborhoods from archaea and bacteria. Genes are shown as arrows from 5’ to 3’. The taxonomic lineage and genomic coordinates for each neighborhood are indicated as are the GenBank genome accessions and, in parentheses, the GenBank accessions for each EVE and DCD protein.

### MmcQ/YjbR-EVE fusion proteins

Related to the RM and TA systems-associated EVE proteins is a class of MmcQ/YjbR-EVE fusions that we found associated with a number of defense gene clusters, as well as signaling, transport, and metabolic factors, mostly, in Firmicutes and γ-proteobacteria (Figure 7). MmcQ/YjbR (PF04237) has a CyaY-like fold and is also fused to tellurite resistance protein TerB and GNAT-type acetyltransferases in other contexts (Anantharaman, Iyer, and Aravind 2012). MmcQ/YjbR-EVE fusions also frequently contain an N-terminal DUF1831 domain, and in many cases, where this domain is missing, there is a DUF1831-MmcQ/YjbR gene immediately adjacent to MmcQ/YjbR-EVE.

DUF1831-MmcQ/YjbR-EVE fusions, which are the most numerous in our data, are frequently encoded within a genomic context that includes sensor histidine kinases, response regulators, and putative DNA-binding proteins. They are also often associated with ABC-type transport system components. Intriguingly, the large number of currently available *Streptococcus* genomes enabled the detection of highly variable regions adjacent to the genes encoding DUF1831-MmcQ/YjbR-EVE proteins in conserved positions. These areas often contain mobile genetic elements (MGEs), defense-associated genes (TA modules, CRISPR-Cas systems), as well as uncharacterized, putative defense, transport, secretory, and DNA/protein repair genes (a MsrAB/disulfide interchange factor operon we detected is likely a mobile protein repair system) (Lourenço Dos Santos, Petropoulos, and Friguet 2018) (Figure 9). These hotspots for integration (and presumably contraction) adjacent to (DUF1831)-MmcQ/YjbR-EVE genes often include transposases, implying a transposon-type mechanism of mobilization. When these variable gene arrays are large, ancestral, independent mobile modules that were assembled to give rise to them can be predicted by comparison with genomes in which the array is smaller (Figure 9). Further work will be necessary to establish the relationship in these systems between the mobile genes and those conserved at the borders, including DUF1831-MmcQ/YjbR-EVE. The fusion of MmcQ/YjbR-EVE to a transposase in *Streptococcus lutetiensis* further underscores that this variety of EVE protein might play a role in regulating the acquisition and/or expression of MGEs. We also observed a similar phenomenon in the regions neighboring MmcQ/YjbR-EVE genes in *Actinobacillus* (Supplementary Figure 4).

### DCD, an EVE-like domain involved in restriction of modified DNA and PCD in plants

We further identified the Development and Cell Death (DCD) domain as a specificity module comparable in sequence and genomic context to EVE. The DCD domain is rare in prokaryotes, and mostly, is present in archaea and hyperthermophilic bacteria. The DCD domain was originally identified in proteins that are strongly induced during plant development, the hypersensitive response to avirulent pathogens, and reaction to various environmental stresses in plants (Tenhaken, Doerks, and Bork 2005; de Camargos et al. 2019; Hoepflinger, Pieslinger, and Tenhaken 2011). Although not classified as such previously, we conclude that DCD is a member of the PUA-like superfamily due to the limited but significant sequence similarity with EVE detected by profile-profile comparison using HHpred (97.16% probability, E-value 0.049). Several of the most highly conserved residues of the EVE domains are present in the DCD domains, and the characteristic secondary structure (βαβαββββ) that forms the EVE β-barrel is also predicted for DCD (Supplementary Figure 9). The DCD domain shows some associations similar to those of defense-related EVE domains, in particular, with Type I restriction systems, as well as a fusion to PD(D/E)XK phosphodiesterases and McrB-like domains, and is distinguished by frequent fusion to a Rossmann-fold methyltransferase, which is extremely rare among EVE domains (Figure 10). These connections imply that, similarly to EVE, DCD domains in prokaryotes recognize methylated bases in DNA and thus contribute to restriction of modified DNA. DCD-methyltransferase fusion protein genes are usually followed by a gene encoding a PD(D/E)XK nuclease, suggesting that they are involved in the additional methylation of modified DNA, recognized by DCD, that could be restricted in the absence of the supplementary methylation.

### DCD and YTH: EVE-like domains with roles in modification-dependent DNA restriction systems and eukaryotic modification-based mRNA processing

We performed a comprehensive search for the DCD domain in all available genomes and found that, among eukaryotes, it is not restricted to plants, as originally described, but is also present in many chromist genomes, particularly, in heterokont and haptophyte algal proteins, where it is often fused to another EVE-like domain, YTH (Figure 11A). The YTH domain is broadly distributed in eukaryotes, has been consistently reported to bind m^6^A in eukaryotic mRNAs, and is involved in multiple processes including splicing and polyadenylation, translation/decay balance (notably triaging of mRNA translation during stress), and inhibition of viral RNA replication (Hazra, Chapat, and Graille 2019; Patil, Pickering, and Jaffrey 2018; Xiao et al. 2016; Liao, Sun, and Xu 2018; Zhao et al. 2019). When fused to the YTH domain in eukaryotes, the DCD domain is also fused to a KH (<u>K h</u>omology) domain, and an array of CCCH-type zinc finger (Znf) domains (Figure 11A). Similar repeated Znfs are conserved in mRNA cleavage and polyadenylation specificity factor 30 (CPSF30) family proteins that are involved in eukaryotic mRNA maturation (Figure 11A) (Valverde, Edwards, and Regan 2008; Shimberg et al. 2016). CPSF30-like proteins in plants also contain a YTH domain and are orthologous to the Znf-Znf-Znf-YTH-DCD-KH proteins we detected. CPSF30 orthologs in fungi and metazoans only have Znf domains, but YTH domain proteins are integral to the CPSF complexes in vertebrates, where they interact with CPSF6 (Figure 11A) (Kasowitz et al. 2018).

**Figure 11:**
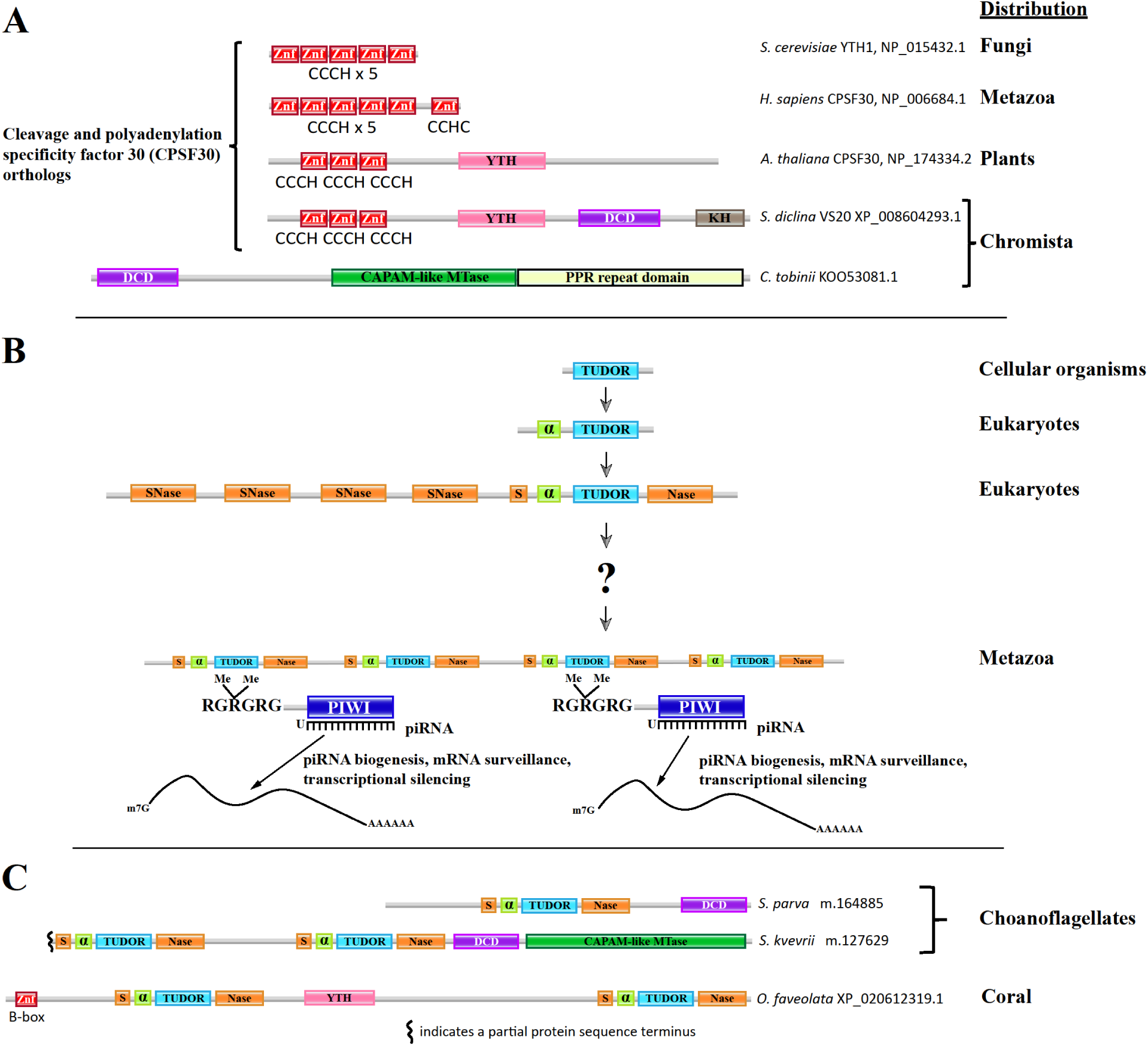
A: Cleavage and polyadenylation specificity factor 30 (CPSF30) orthologs contain YTH and DCD domains in many heterokont species, and a CAPAM-like putative mRNA methyltransferase containing a DCD domain was detected in a haptophyte species; B: Multi-eTudor domain proteins evolved from Tudor-SN to produce the family of essential piRNA pathway factors in metazoans, but their ancestry is poorly understood; C: eTudor-DCD proteins in choanoflagellates and eTudor-YTH proteins in corals clarify the evolutionary history of the eTudor family. A: Domain organization of representative orthologs of CPSF30 from budding yeast, human, *Arabidopsis*, and the pathogenic oomycete *S. diclina*, as well as a CAPAM-like putative mRNA methyltransferase from *C. tobinii* which contains a DCD domain and was the only example detected. The RefSeq protein accession numbers are indicated. Zinc finger (Znf) domains are labelled with their type. B: The core Tudor domain inherited from prokaryotes and present in histone-binding proteins is shown in blue. An N-terminal α-helix conserved in SMN-like proteins and incorporated into the eTudor domain is shown in green. The SNase domain that the SMN-like Tudor domain inserted into is shown in orange, with sections at both termini of the resulting eTudor domain that fold around the inserted Tudor domain to retain the original SNase structure (Liu et al. 2010). A multi-eTudor protein is shown interacting with a dimethylarginine residue in an RGRGRG motif at the N-termini of two Piwi-related Argonautes. Such motifs are conserved sites of arginine methylation in the Argonautes, as well as Sm proteins bound by the Tudor domains in SMN and Tudor-SN. C: Domain organization of two choanoflagellate eTudor-DCD proteins and a representative coral eTudor-YTH protein. The species and GenBank or RefSeq protein accession numbers, or identifiers from the predicted protein dataset published by Richter et al. are indicated (Full sequences in Supplementary data). 5’ partial proteins predicted from transcriptome sequencing are marked at the incomplete terminus. Zinc finger (Znf) domains are labelled with their type.

YTH also has been reported to bind m^6^A in DNA when fused to an McrB homolog in the archaeon *Thermococcus gammatolerans* (Hosford, Bui, and Chappie 2020). Using the sequence of this archaeal YTH domain as a PSI-BLAST query, we detected homologs that, much like DCD, are fused to McrB-like GTPases or PD(D/E)XK nucleases. Most of these YTH-like domains are not clearly distinguishable from EVE domains using HHpred, being modest hits for both types, a pattern that is reminiscent of some prokaryotic DCD domains.

### Extended Tudor-DCD fusion proteins in choanoflagellates implicated in the origins of the piRNA pathway

We were unable to identify DCD domains in metazoans. However, when we analyzed the predicted proteins translated from the published transcriptomes of choanoflagellates, the closest unicellular relatives of animals (Hoffmeyer and Burkhardt 2016; Richter et al. 2018), a protein containing a DCD domain fused to an extended Tudor (eTudor) domain was detected in both loricate and non-loricate choanoflagellates, the two main lineages of this phylum (Figure 11C, 13).

Tudor domains bind post-translationally methylated arginine or lysine residues in eukaryotic proteins (Chen et al. 2011). They interact with three main types of modified proteins: histone tails (methylarginine or methyl-lysine), Sm proteins in spliceosomes (methylarginine), and the N-termini of metazoan Piwi-related Argonaute proteins (methylarginine) (Chen et al. 2011). The Sm protein-binding Tudor domains present in the splicing factor Survival Motor Neuron (SMN) and related proteins are distinguished from Tudor domains that bind histone tails by an N-terminal α-helix (Figure 11B) (Jin et al. 2009). The Tudor domains that interact with Piwi-related Argonautes are of the eTudor type (Chen et al. 2011; Liu et al. 2010). The eTudor family is restricted to metazoans, with the exception of Tudor-SN, a highly conserved eukaryotic protein implicated in RNA interference, splicing, microRNA decay, and RNA editing that contains four staphylococcal nuclease (SNase) domains and a single eTudor domain (Li et al. 2008; Gao et al. 2012; Elbarbary et al. 2017) (Figure 11B).

Bioinformatic and structural analyses suggest that the eTudor domain arose when a Tudor domain, related to the Tudor domain in SMN, inserted into the fifth, C-terminal SNase domain of an ancestral multi-SNase protein (Jin et al. 2009; Liu et al. 2010) (Figure 11B). The resulting domain fusion of Tudor and SNase (hence the name ‘extended Tudor’), became the ancestor of the eTudor family, in which the catalytic residues from the ancestral SNase domain are mutated, likely, rendering it inactive (Jin et al. 2009; Liu et al. 2010; Li et al. 2008; Li et al. 2018) (Figure 11B). Present in all metazoans, multi-eTudor proteins play crucial roles in the localization of Piwi-related Argonautes and biogenesis of Piwi-interacting RNAs (piRNAs) by interacting with symmetrically dimethylated arginine (SDMA) residues in the Argonaute N-termini, and thus, are essential for repression of transposable elements, modulation of germline mRNA levels, and germ/stem cell immortality (Tóth et al. 2016; Sturm et al. 2017; Chen et al. 2011). The origins of the complex metazoan multi-eTudor proteins derived from Tudor-SN are fundamental to the understanding of the piRNA pathway and animal germline specification but, currently, remain obscure.

We detected eTudor proteins in choanoflagellates that are not orthologs of Tudor-SN, and these represent the first examples, to our knowledge, to be reported in a non-metazoan organism. These proteins usually also contain a DCD domain, and in some cases, a CAPAM (<u>cap</u>-specific <u>a</u>denosine <u>m</u>ethyltranferase)-like methyltransferase and/or a second eTudor domain (Figure 11C, 13). In the process of identifying the CAPAM-like domains, we encountered a misannotation of the Pfam family PCIF1_WW (pfam12237), which, according to our analysis, is not a WW domain, but rather, a CAPAM-like methyltransferase. We also observed multi-eTudor proteins fused with a YTH domain in some species of coral, among the earliest branching metazoans (Figure 11C, 13).

We also detected links between the eTudor-DCD/YTH fusion proteins and the ubiquitination pathway and protein degradation. An N-terminal ubiquitin-binding domain (UBA) and B-box Znf domains are present in the choanoflagellate eTudor-DCD and coral eTudor-YTH proteins, respectively (Figure 13). Similar B-box Znfs are found in TRIM ubiquitin E3 ligases and in the eTudor piRNA pathway factor qin/komo from *Drosophila*, which also contain RING Znfs (Figure 13) (Chen et al. 2011; Tomar and Singh 2015). The eTudor proteins with RING Znfs are conserved throughout the eumetazoa, although in vertebrates, the B-box Znfs appear to have been lost. In the sponge *Amphimedon queenslandica*, a protein with four eTudor domains and an N-terminal MYND-type Znf has been identified, with orthologs present in most metazoans (Figure 13).

**Figure 12:**
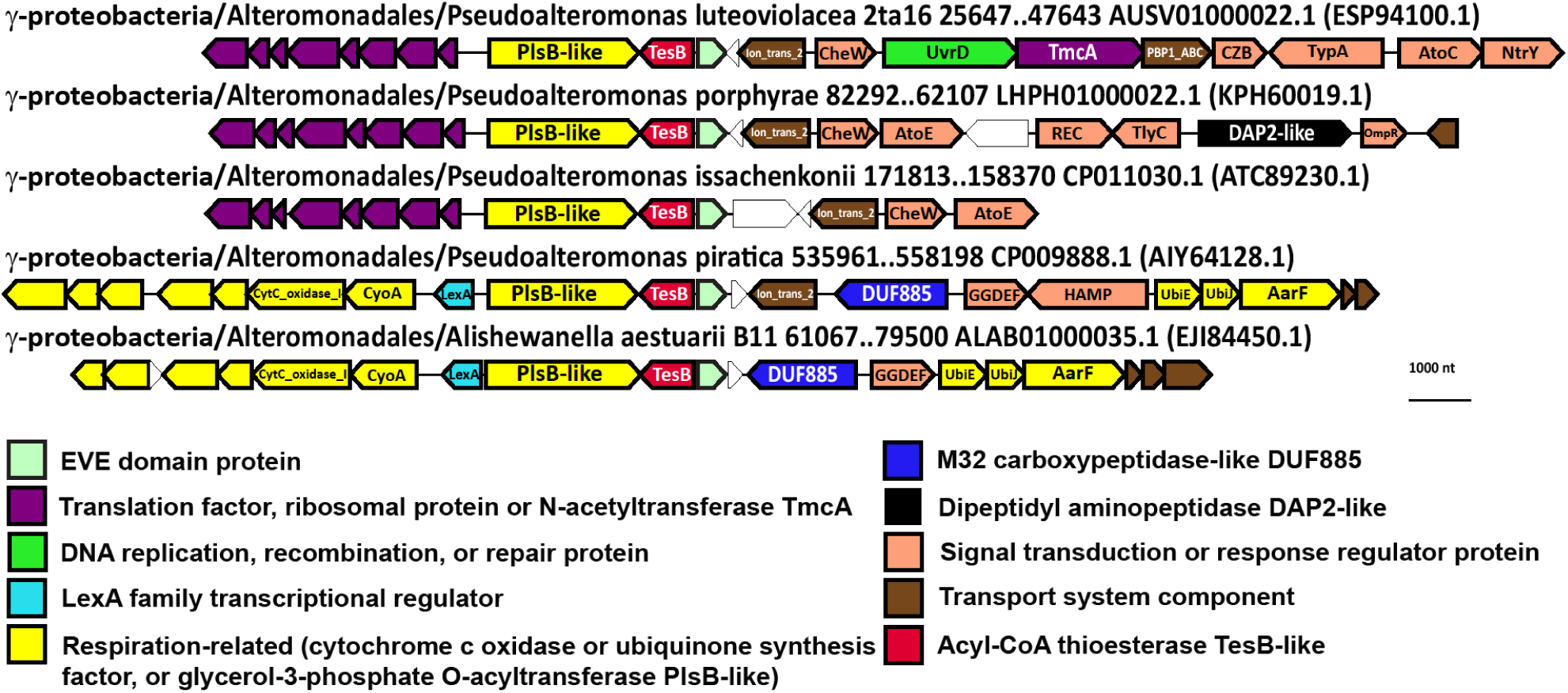
Conserved genomic context of EVE proteins in Pseudoalteromonadales. Representative EVE protein neighborhoods from Pseudoalteromonadales. Genes are shown as arrows from 5’ to 3’. The order of α-proteobacteria, species, and genomic coordinates for each neighborhood are indicated, as are the GenBank genome accessions and, in parentheses, the GenBank accessions for each EVE protein.

**Figure 13:**
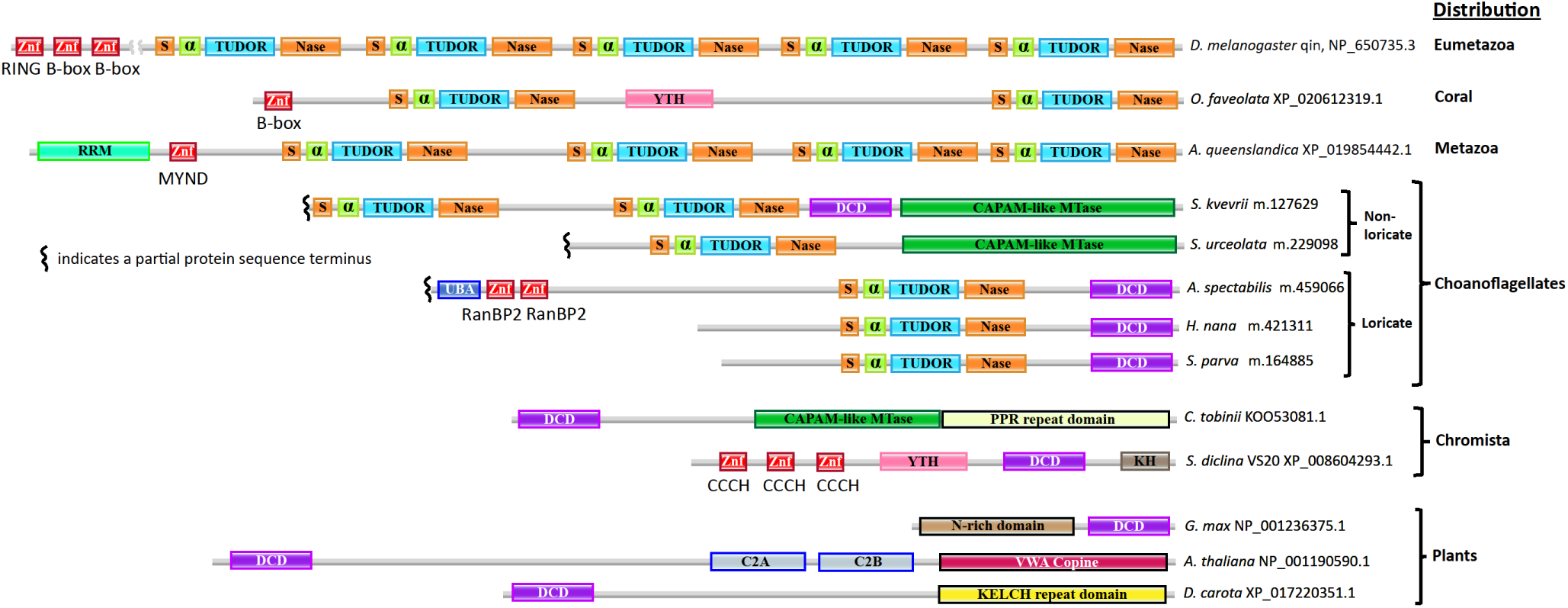
Representatives of major classes and informative examples of DCD and eTudor proteins in plants, chromists, choanoflagellates, and metazoans. Domain organization of representatives of major classes of DCD proteins in plants and heterokonts, and major classes of coral and eumetazoan eTudor proteins. Especially informative individual examples from haptophyte algae and choanoflagellates are also shown. The species and GenBank or RefSeq protein accession numbers, or identifiers from the predicted protein dataset published by Richter et al. are indicated (Full sequences in Supplementary data). 5’ partial proteins predicted from transcriptome sequencing are marked at the incomplete terminus. Zinc finger (Znf) domains are labelled with their type. The phyletic distribution of each protein class is denoted as well. Choanoflagellate protein sequences that are not deposited as separate entries in GenBank are available as Supplementary Dataset 2.

### Biochemical functions of EVE-associated proteins

In this section, we present our inferences of the likely biochemical functions of the proteins linked to the EVE domain, both covalently and non-covalently, which we derived from the literature documenting experimental characterization of members of the corresponding protein families. An important caveat is that these inferences, although often direct and likely valid, are inherently less confident than the robust computational results so far described.

#### EVE in α-proteobacteria

The most prominent contextual association of the EVE domain observed in our study is its inclusion in the putative operon TsaD->GpsA->YciI->EVE in α-proteobacteria. TsaD modifies tRNAs that decode ANN codons (Met, Ile, Thr, Asn, Lys, Arg, Ser) to introduce threonylcarbamoyladenosine (t^6^A) at position 37, immediately adjacent to the anticodon (Swinehart et al. 2020). t^6^A is a universal modification that is essential for translational fidelity, and mutations in this pathway lead to errors in start codon selection and aberrant frameshifts (Missoury et al. 2018; Swinehart et al. 2020; Deutsch et al. 2012). The α-proteobacterial glycerol-3-phosphate dehydrogenase that is tightly associated with EVE is orthologous to the corresponding mitochondrial enzyme that contributes electrons to the respiratory chain (Mráček, Drahota, and Houštěk 2013). Thus, the evolutionarily conserved link between the EVE proteins, TsaD and GpsA implies an unexplored connection between tRNA modification and electron transfer in α-proteobacteria. The YciI protein family has not been thoroughly characterized. One member of this family, TftG from *Burkholderia phenoliruptrix* AC1100, is a dehydrochlorinase requiring a conserved His-Asp dyad for catalysis, a motif that is present in the YciI proteins in the EVE neighborhoods (Hayes et al. 2013). Fusion of a YciI domain to a s^70^ factor domain in *Caulobacter vibrioides* and to a BolA transcriptional regulator domain in *Coxiella burnetii* imply that this family may be involved in transcription initiation (Dressaire et al. 2015). Intriguingly, in *E. coli* K-12, the gene endoding TsaD is in a head to head orientation with an operon that encodes σ^70^ factor RpoD, suggesting that a link between tRNA modification and global transcriptional regulation could be ancestral to Proteobacteria. This apparent operon is frequently associated, in a head to head orientation implying possible co-regulation, with a HemC->HemD->IMMP->HemY operon that encodes enzymes of heme biosynthesis. The IMMP ortholog in mitochondria is required for the formation of cristae (von der Malsburg et al. 2011).

#### EVE in β and γ-proteobacteria

The cell division proteins strongly associated with EVE in β and γ-proteobacteria, ZapA and ZapB, interact as a complex with FtsZ, promoting the Z-ring formation during bacterial cytokinesis (Galli and Gerdes 2010). Often located between ZapAB and EVE is the enzyme FAU1 (or YgfA), which converts 5-formyltetrahydrofolate (5-formylTHF) to 5,10-methylenylTHF. 5-formylTHF is a stable storage form of folate that accumulates in dormant cells, such as spores and seeds, whereas 5,10-methylenylTHF is a precursor in purine and methionine biosynthetic pathways (Field, Szebenyi, and Stover 2006; Kruschwitz et al. 1994; Stover and Schirch 1993). Also present in these putative operons, always immediately following ZapA genes, are non-coding 6S RNA (*ssrS*) genes, which express a 184 nucleotide small RNA that functions as a global regulator of transcription in bacteria by binding to the housekeeping σ^70^-RNA polymerase holoenzyme (Eσ^70^) (Wassarman and Storz 2000; Wassarman 2018). 6S RNAs have been reported to accumulate in *E. coli* cultures during the transition from the exponential to stationary phase of growth, and their effect is to inhibit transcription from most σ^70^ -dependent promoters, which effectively activates the expression of stationary phase-specific genes dependent upon other σ factors, enabling transcriptional adaptation to changing growth conditions (Wassarman and Storz 2000; Steuten et al. 2014).

It appears likely that expression of FAU1, in conjunction with ZapAB, SsrS, and the EVE protein, is part of a metabolic switch between proliferative modes. Consistent with this possibility, FAU1 has been implicated in promoting the formation of persister cells and biofilms, and SsrS function is thought to enhance long term cell survival (Trotochaud and Wassarman 2004; Hansen, Lewis, and Vulić 2008; Ren et al. 2004). The SsrS->FAU1 operon, which is broadly conserved in Proteobacteria, including α-proteobacteria, has been experimentally characterized in *E. coli* K-12, where the dicistronic transcript is processed into mature 6S RNA (Chae et al. 2011; Kim and Lee 2004). Our observations suggest that FAU1 genes are not strictly necessary in these regions, and that *ssrS* genes are often flanked, in β and γ-proteobacteria, but not in α-proteobacteria, by ZapA and EVE genes, which may, like FAU1, be expressed in polycistronic transcripts containing the 6S RNA precursor. No modifications of 6S RNA have been reported although the consistent, close association with EVE suggests that the 6S RNA might contain modified bases recognized by EVE domains (Figure 5).

Given its conserved, head to head juxtaposition in γ-proteobacteria with the apparent ZapAB->SsrS->(FAU1)->EVE operon, the operon YgfB->PepP->UbiH->UbiI that has been experimentally characterized in *E. coli* K-12 (Nakahigashi et al. 1992), could be co-expressed and might play a role in cell cycle regulation as well. The PepP ortholog encoded by *Pseudomonas aeruginosa* in this conserved, EVE-containing context has been identified as a critical virulence factor in a *Caenorhabditis elegans* infection model (Feinbaum et al. 2012). Homologs of ribose-5 phosphate isomerase (RpiA) and L-threonine dehydratase (IlvA) that are often encoded in these regions likely also participate in the implied, large-scale proliferative regulation.

Alteromonadales, while lacking the link between EVE and ZapAB->SsrS, encode a cobalamin-dependent radical SAM enzyme in close association with EVE, which might be related to the cobalamin biosynthetic clusters adjacent to ZapAB->SsrS->EVE in β-proteobacteria (Figure 5, Supplementary Figure 3). Pseudoalteromonadales, also lacking ZapAB->SsrS->EVE, encode EVE domains in conserved associations with translation factors and respiration-related enzymes involved in the maturation of cytochrome c and ubiquinone, suggesting that coupling between translation and respiration mediated by EVE domains extends throughout the Proteobacteria and could be ancestral to this phylum (Figure 12). In these neighborhoods, EVE is tightly linked to factors homologous to acyl-CoA thioesterase TesB and glycerol-3-phosphate O-acyltransferase PlsB, suggesting that the abundance of glycerol-3-phosphate, which can contribute electrons to the respiratory chain via its dehydrogenase (Mrácek, Drahota, and Houštěk 2013), as seen in the α-proteobacterial EVE neighborhoods, is modulated by these EVE-associated enzymes.

Our comparative genomic analyses shed light on deeply conserved apparent functions of EVE proteins in α-proteobacteria, where they likely link modulation of cytochrome c maturation with tRNA modification and transcriptional regulation, and in β and γ-proteobacteria, where they are predominantly implicated in the linkage of cell division, transcriptional, and metabolic regulatory mechanisms, but in some members of these classes, are closely associated with translation and electron transport factors as in α-proteobacteria. As noted above, in *Coxiella burnetii*, a YciI-like protein is fused at the C-terminus to a BolA domain, a transcriptional regulator involved in promoting biofilm formation and repressing motility (Dressaire et al. 2015; Willis et al. 2005). Morphological effects of BolA overexpression depend on FtsZ, the interaction partner of ZapAB (Aldea et al. 1988). This finding suggests that YciI proteins closely tied to EVE in α-proteobacteria might perform a role similar to the YciI homolog fused to BolA, which is likely to involve transcriptional regulation of large-scale biochemical and morphological adaptations to changing conditions. A similar function is conceivably carried out by the ZapAB->SsrS->(FAU1)->EVE regions in β and γ-proteobacteria, which likely participate in cellular phase shifts between exponential vs. stationary and planktonic vs. biofilm proliferative modes.

#### EVE as a specificity domain in modification-dependent restriction systems

The most common function of EVE domains in this capacity likely entails flipping out a modified cytosine derivative from a DNA helix for scrutiny in the EVE’s binding pocket and targeting endonuclease activity to the neighboring DNA, given sufficient affinity for the modified sequence. This role can be inferred from the comparison with the SRA domain, which shares the PUA-like fold with EVE. The SRA domain has been characterized in considerable detail, including the base-flipping 5mC DNA-binding mechanism and characterization of its function as a modified DNA specificity module in Type IV REs which restrict DNA containing 5mC, 5hmC, and glucosylated 5hmC (Arita et al. 2008; Avvakumov et al. 2008; Hashimoto et al. 2008; Liu et al. 2018; Weigele and Raleigh 2016). Therefore, EVE domains deployed in modification-dependent restriction likely use a base-flipping mechanism, similar to that of SRA, to sense modified cytosine in various sequence contexts although some might bind derivatives of the other pyrimidine bases, thymine or uracil, which are hypermodified in some phage genomes (Weigele and Raleigh 2016). Yet other EVE domains might preferentially bind modified adenine, given that EVE is also structurally similar to the YTH domain, which binds m^6^A in DNA and RNA (Bertonati et al. 2009; Hosford, Bui, and Chappie 2020).

We detected a remarkable variety of combinations of EVE and restriction endonuclease domains, implying intense pressure to evolve diverse restriction strategies to provide immunity from a vast and highly varied population of viruses with modified genomes. Some of these modifications bound by EVE domains could be effective in inhibiting defense by CRISPR-Cas adaptive immunity systems, in addition to Type I, II, and III REs, as has been reported for glucosylated 5hmC (Vlot et al. 2018; Weigele and Raleigh 2016). The EVE domains appear to play a major role in meeting the demand for defenses tailored to this threat.

#### EVE domains in toxin-antitoxin systems

None of the modules (PIN, GNAT, HTH, EVE, ASCH) in these systems have been experimentally characterized, but the predicted toxin or antitoxin activity of the associated domains strongly suggests that, in this case, the EVE proteins are components of toxin-antitoxin (TA) systems. The PIN domain RNases function as toxins in a broad variety of bacterial, and especially, archaeal TA systems (Matelska, Steczkiewicz, and Ginalski 2017; Makarova, Wolf, and Koonin 2009). The GNAT domains also typically function as toxins (Yeo 2018), and HTH domain-containing antitoxins have been described as well (Chan, Espinosa, and Yeo 2016). The mechanistic details of these (PIN)-GNAT-EVE protein activities await experimental investigation, but some functional hints emerged from our analysis. The differential distributions of PIN-GNAT-EVE and GNAT-EVE proteins (Figure 8) is a potentially important clue. Furthermore, in *Methanohalophilus* genomes, the associated ASCH domain is fused to the specificity subunit (HsdS) of a Type I RM system, implying that it confers modification specificity to the RM complex (Figure 8). Combined with the frequent occurrence of HTH-ASCH fusions, these associations suggest that ASCH domains in the (PIN)-GNAT-EVE operons bind modified DNA.

The uncharacterized, putative defense system we detected, COG1743->DUF499->SWI2/SNF2-nuclease->EVE, and the Type I RM systems associated with standalone EVE and ASCH proteins, arguably represent TA systems as well (Supplementary Figure 8, Figure 10). We predict that DUF499 proteins in the former system might interfere with the replication of foreign DNA containing modified bases, which is discriminated by the EVE domain and restricted by the SWI2/SNF2 helicase-nuclease fusion protein, activities that are potentially toxic. The likely role of the methyltransferase is to prevent restriction of the host genome, and thus to serve as an antitoxin, by methylating a base in the sequence recognized by the SWI2/SNF2-nuclease, a modification not recognized by the EVE domain. The presence of this factor also implies that the system can restrict unmodified DNA without requiring recognition by the associated EVE domain.

Similarly, Type I RM systems that are associated with EVE domains are likely to target modified DNA, and the presence of the Type I methyltransferase (HsdM) suggests restriction of both unmodified and modified DNA can occur (Figure 10). The methyltransferase in these systems can be predicted to generate modified bases that are not recognized by the associated EVE domain and prevent restriction by the Type I endonuclease subunit (HsdR), which is also capable of restricting modified DNA that is discriminated by the EVE domain.

#### MmcQ/YjbR-EVE fusion proteins

The homology between MmcQ/YjbR and the mitochondrial iron homeostasis protein CyaY suggests that MmcQ/YjbR could be an iron-binding protein as well (Layer et al. 2006) although the functional residues and electrostatic potential are not conserved between these domains (Singarapu et al. 2007). A more convincing functional prediction for MmcQ/YjbR has been made based on structural and electrostatic surface similarity to the C-terminus of T4 bacteriophage transcription factor MotA, known as MotCF. Although there is only a limited sequence similarity between with MmcQ/YjbR and MotCF, conserved residues are concentrated in the putative DNA-binding region of MotCF, strongly suggesting that, like MotCF, MmcQ/YjbR interacts with DNA (Singarapu et al. 2007). Furthermore, multiple MmcQ/YjbR homologs have been shown to adopt a ‘double wing’ DNA-binding fold similar to MotCF (Singarapu et al. 2007; Feldmann et al. 2012). In our dataset, one example of a GIY-YIG nuclease-MmcQ/YjbR-EVE fusion and another of a PLDc nuclease-Helicase-MmcQ/YjbR-EVE fusion are present, implicating the EVE domain as a modified DNA base specificity module in these proteins, whereas MmcQ/YjbR might contribute sequence specificity.

DUF1831, often fused at the N-terminus to MmcQ/YjbR-EVE proteins shows remote structural similarity to TBP-like (TATA-binding) fold proteins, which include S-adenosyl-methionine decarboxylase (Bakolitsa et al. 2010). Analysis of the genomic neighborhood context of DUF1831 genes supports a role in metabolism of amino acids, particularly, methionine (Bakolitsa et al. 2010). The DUF1831-MmcQ/YjbR-EVE proteins can be encoded in a putative operon with peptide methionine sulfoxide reductase MsrAB and disulfide interchange factors which likely recycle it. In general, however, the function of these complex EVE proteins is likely to be multifaceted (Figure 9).

#### Extended Tudor-DCD fusion proteins in choanoflagellates implicated in the origins of the piRNA pathway

The identification of eTudor-DCD fusion proteins encoded in the transcriptomes of loricate and non-loricate choanoflagellates implies that the common ancestor of the extant choanoflagellates as well as the metazoa, that apparently descended from colonies of non-loricate choanoflagellate-like ancestors (Hoffmeyer and Burkhardt 2016; Cavalier-Smith 2017), already encoded the eTudor-DCD protein. Choanoflagellates, most likely, acquired the genes encoding DCD proteins by horizontal gene transfer (HGT) following the loss of DCD in the ancestor of the opisthokonts. This route of evolution is strongly suggested by the observed absence of DCD from all opisthokonts, other than the choanoflagellates, although the possibility of inheritance from an early eukaryotic ancestor cannot be completely ruled out (Tenhaken, Doerks, and Bork 2005). Consistent with this scenario, the extant choanoflagellates are thought to have acquired substantial portions of their genomes via HGT, including from algae (Tucker 2013; Yue et al. 2013). In both haptophyte algae (chromists) and non-loricate choanoflagellates, the DCD domain is fused to a CAPAM-like domain, and in the latter case, also to an eTudor domain (Figure 13) (Akichika et al. 2019). Furthermore, as noted above, DCD is present in conserved CPSF30-like factors in many heterokont species (also chromists). Therefore, we infer that chromist algae are the putative source of DCD in choanoflagellates that might have acquired it via HGT.

In heterokont DCD proteins that are orthologs of CPSF30 and so can be predicted to participate in mRNA maturation, the DCD domain likely binds modified RNA, either exclusively or in addition to binding modified DNA. Its primary target could be m^5^C in mRNA, given the affinity of EVE domains for modified cytosine, and the presence, in the same proteins, of a YTH domain, which has consistently been reported to bind m^6^A in eukaryotic mRNA (Anders et al. 2018; Hazra, Chapat, and Graille 2019; Kasowitz et al. 2018; Xiao et al. 2016; Patil, Pickering, and Jaffrey 2018; Zhao et al. 2019; Liao, Sun, and Xu 2018). It is probable that the DCD and YTH domains in these proteins recognize distinct modifications. It is this type of DCD domain, which is likely to bind modified RNA, that can be predicted to have fused to eTudor in choanoflagellates.

The nature of the ligand of the choanoflagellate DCD domains is suggested by the other domains to which they are covalently linked. The CAPAM-like methyltransferase fused to DCD in *S. kvevrii* (a choanoflagellate) and *C. tobinii* (a haptophye alga) is homologous to the RNA methyltransferase in the CAPAM protein which methylates m^6^A in vertebrate mRNAs (Figure 13) (Akichika et al. 2019). In the DCD-CAPAM-like methyltransferase fusion proteins, DCD occupies the same N-terminal position as a helical domain that, in CAPAM, is involved in the recognition of the m^7^G mRNA cap and directing methylation to m^6^A (Akichika et al. 2019). Therefore, the DCD domain in these proteins might be involved in targeting additional modifications of modified mRNAs, perhaps, containing m^5^C. Furthermore, the presence of N-terminal RanBP2-type Znfs in the eTudor-DCD protein we detected in *A. spectabilis* also implies RNA binding, as well as participation in splicing and/or nuclear export (Figure 13) (Nguyen et al. 2011; Ritterhoff et al. 2016). The eTudor domain itself, in the absence of Piwi-related Argonaute proteins, is likely to bind SDMA residues in spliceosomal Sm proteins during mRNA maturation, as shown for the homologous domains in the Tudor-SN and SMN proteins (Gao et al. 2012; Selenko et al. 2001).

Similar to the choanoflagellate eTudor-DCD protein we identified in *A. spectabilis*, which contains an N-terminal UBA domain, metazoan eTudor proteins often contain N-terminal Znfs implicated in ubiquitination and protein degradation, and one well-conserved type possesses N-terminal MYND-type Znfs (Figure 13). The MYND-type Znf in the *Aedes aegypti* eTudor protein Veneno, which contains two eTudor domains, is required for the localization of Veneno to putative piRNA processing germ granules (Joosten et al. 2018). The consistent presence of N-terminal Znf domains, in many metazoan eTudor proteins and one loricate choanoflagellate eTudor protein, led us to surmise that the incomplete N-termini in the partial eTudor protein sequences we identified in non-loricate choanoflagellates likely harbor a type of N-terminal Znf as well, which has not yet been observed (Figure 13, 14).

**Figure 14:**
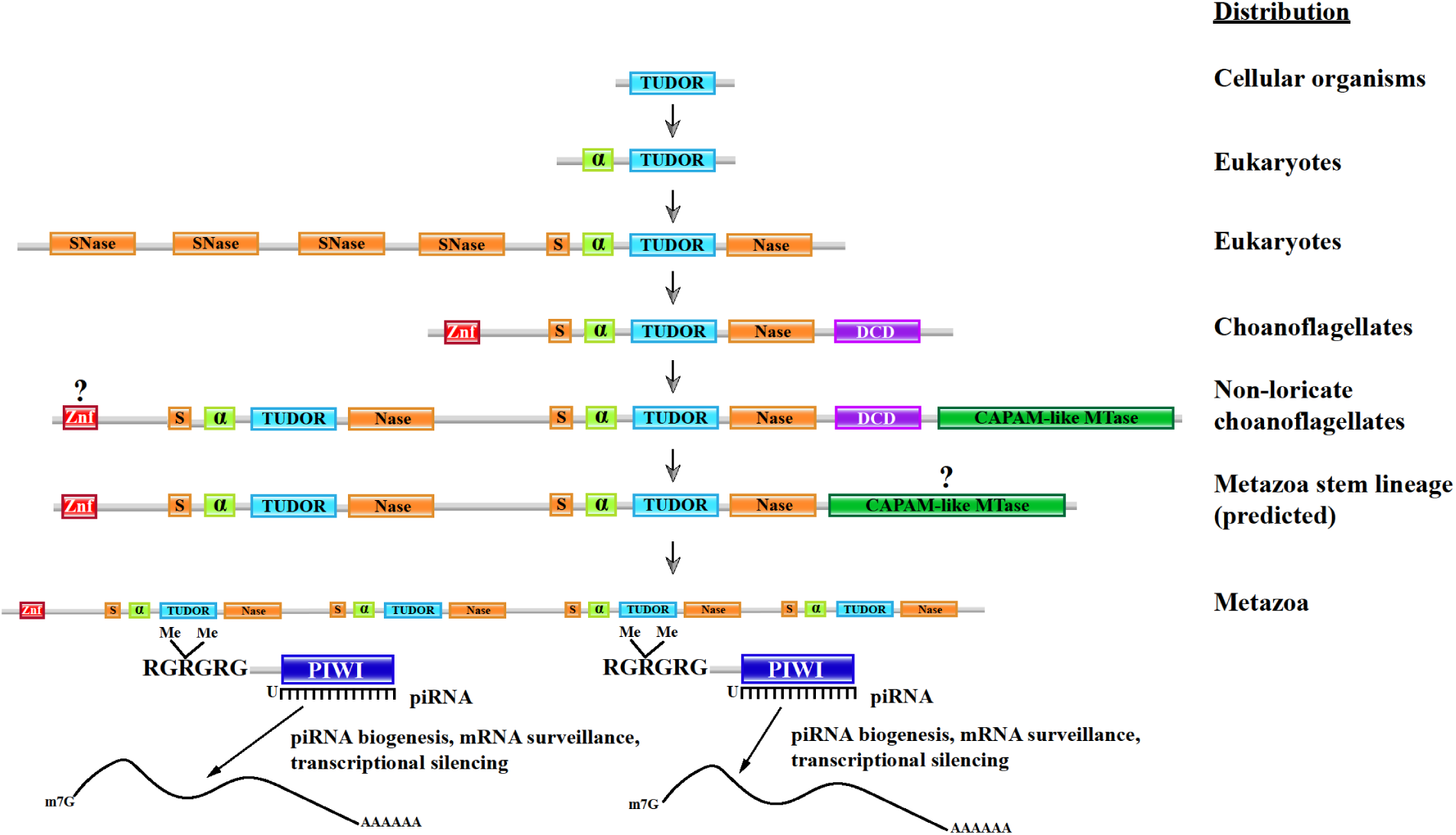
A hypothetical evolutionary scenario for the metazoan eTudor domain proteins essential to the piRNA pathway. The predicted evolution of Tudor protein function and domain composition is depicted. In the first stage, the core Tudor domain is involved in recognition of methylarginine in histone tails, with an unknown prokaryotic role. Next, this domain is combined with an N-terminal α-helix in an ancestral SMN-like splicing factor in the eukaryotic stem lineage, binding SDMA in spliceosomal Sm proteins. Subsequently, but also during the eukaryotic stem phase, this type of Tudor domain was inserted into an SNase domain in the ancestor of the Tudor-SN-like family, forming the eTudor domain. The Tudor-SN protein class became involved in RNAi and splicing, binding SDMA in spliceosomal Sm proteins. During the diversification of eukaryotes, in ancestral choanoflagellates, the eTudor domain was fused to the DCD domain, forming a putative mRNA maturation factor binding SDMA in spliceosomal Sm proteins with eTudor and m^5^C RNA with DCD. This type of protein may play a role in mRNA nuclear export, self vs. non-self discrimination, and protein degradation, with N-terminal Znf domains that might bind ubiquitin or RNA. At the next step, after the divergence of the loricate and non-loricate choanoflagellates, duplication of the eTudor domain in the non-loricate choanoflagellates allowed elaboration of the function of the eTudor-DCD protein, freeing one eTudor domain from the conserved functional requirement of Sm protein binding and allowing it to interact with other proteins containing SDMA, eventually including Piwi-related Argonautes, but perhaps not yet at this stage. The involvement of the CAPAM-like methyltransferase is not necessarily part of the piRNA pathway origins, but the possibility is suggested by the data. In the final link to the extant piRNA systems, we propose a hypothetical factor, potentially still binding Sm proteins, that evolved to target mRNA via interaction with piRNA/Argonaute complexes rather than binding m^5^C. These complexes now discriminate spliced and modified self from non-self/aberrant mRNA, regulating nuclear export, as well as transcription via Argonaute directed chromatin modification, a conserved feature of eukaryotic cells also extensively performed by plants. N-terminal Znfs contribute to eTudor localization to perinuclear germ granules and possibly link them to the ubiquitination and protein degradation system to provide even more control over expression.

## Discussion

The comprehensive analysis of the genomic neighborhoods of prokaryotic EVE proteins described here has a variety of functional and evolutionary implications.

### Implications for the evolution of PCD from proteobacterial and defense-related EVEs

PCD in eukaryotes reportedly involves THYN1-like EVE domains, which are broadly distributed and show high sequence similarity to proteobacterial EVEs. The conservation of the EVE genomic context in α-proteobacteria and Pseudoalteromonadales suggests the possibility that eukaryotic PCD evolution exploited a proteobacterial mechanism that couples modulation of energy production with translation via cytosine methylation in tRNAs, along with the EVE proteins that recognize these modifications. Intriguingly, roles for tRNA and stress-induced, tRNA-derived tiRNAs in the intrinsic PCD pathway have recently come to light. Multiple studies have demonstrated an interaction between tRNA/tiRNAs and cytochrome c in mammalian cells that inhibits the formation of the apoptosome and promotes cell survival (Mei et al. 2010; Saikia et al. 2014). In the case of tiRNAs, which are generated from tRNA cleavage near the anticodon, modifications of the tRNA, such as 5-methylcytosine (m^5^C), which might be recognized by an EVE domain, have been reported to negatively regulate their biogenesis (Saikia and Hatzoglou 2015). Furthermore, mitochondrial IMMP, which is orthologous to the protein closely associated with the EVE domain in α-proteobacteria, has been implicated in eukaryotic PCD (Madungwe et al. 2018). In addition, the heme biosynthesis enzymes encoded in the same neighborhoods with EVE proteins in α-proteobacterial are involved in the maturation of cytochrome c, which requires heme as a cofactor, and therefore, are linked to one of the central effectors of eukaryotic PCD (Kranz et al. 2009).

Moreover, the neighborhoods of EVE proteins in β and γ-proteobacteria implicate the EVE domain in deeply conserved coordination between proteins that promote cytokinesis (ZapAB), a small RNA that promotes transcriptional adaptation to growth conditions (SsrS, which may be modified and bound by an EVE domain), and a metabolic enzyme (FAU1) involved in persister cell and biofilm formation under environmental stress. In γ-proteobacteria, an associated operon encoding a virulence-related aminopeptidase (PepP) and respiration-related factors of ubiquinone biosynthesis (UbiHI) is also likely to contribute to the overall function of these conserved regions. We cannot yet determine the ancestry of the eukaryotic EVE domains, but these roles of EVE proteins in bacteria tying together energetic, transcriptional, and translational responses could presage the involvement of this domain in eukaryotic PCD.

Defense-related EVE domains, which do not generally fall into the two largest clusters from our CLANS analysis, nevertheless, are likely to be involved in PCD. Modification-dependent restriction systems are generally toxic to cells which express enzymes that catalyze the formation of modified bases they recognize, and thus, are implicated in a form of prokaryotic PCD (Weigele and Raleigh 2016).

Therefore, EVE domains associated with nucleases are potentially involved in both innate immunity and PCD, in cases when a cognate methyltransferase is expressed in the same cell. The connection between TA systems containing EVE domains and its role in prokaryotic PCD is readily apparent. The available evidence concerning MmcQ/YjbR-EVE fusion proteins suggests coordination of environmental sensing and response, metabolism, and defense/PCD that is modulated by that class, which represents yet another way in which EVE domains participate in the complex chains of events involved in cell fate decisions. The DCD domain, a defense-related EVE-like domain, likely shares the innate immunity/PCD role of EVE in prokaryotes, whereas in eukaryotes, it has taken on more complex functions that could involve modified RNA binding and has resulted clear involvement of this domain in PCD in plants, as well as a likely role in mRNA maturation in chromists and choanoflagellates, which was apparently important during the evolution of the eTudor proteins and the piRNA pathway in metazoans.

### Involvement of (PIN)-GNAT-EVE proteins in virus-host conflicts

In light of the observations described above concerning (PIN)-GNAT-EVE proteins, targeting GNAT to modified DNA or RNA via EVE, conceivably, protects phages against host RM systems, perhaps, via toxicity to the host, whereas the addition of PIN is associated with host defense and likely counteracts the effect of GNAT-EVE. The GNAT-EVE proteins potentially target aminoacyl-tRNAs (aa-tRNAs) that harbor modifications recognized by EVE, given that GNAT toxins have been reported to acetylate aa-tRNAs (Wilcox et al. 2018; Yeo 2018). Under this scenario, PIN-GNAT-EVE proteins would likely degrade toxic, acetylated aa-tRNAs generated by GNAT-EVE. The role of the putative DNA-binding ASCH domain that is nearly always present in (PIN)-GNAT-EVE operons in this process remains unclear, but it might be involved in regulating the expression of the (PIN)-GNAT-EVE protein. Other accessory proteins, such as AAA_17 family ATPases that are frequently, but not invariably, encoded near these factors can be expected to contribute in non-essential, regulatory capacities. The interplay between phage and host cell proteins with GNAT-EVE architectures appears to be a widespread phenomenon that clearly warrants further inquiry.

### Extended Tudor-DCD and the origins of the piRNA pathway

We predicted that DCD proteins in choanoflagellates and chromista bind modified RNA, or less likely, also DNA. It is unclear whether DCD domain proteins in plants, which have been the subject of considerable inquiry, bind modified DNA, RNA, or both, but investigation of their affinities toward modified nucleic acids can be expected to elucidate their roles in plant PCD. Conceivably, given the distributions we observed and the domains fused to DCD and YTH in various phyla, these domains originate from fast-evolving EVE domains involved in restriction of modified DNA that were recruited, early in eukaryogenesis, to recognize modified eukaryotic mRNA.

Mechanistically, the eTudor-DCD proteins in choanoflagellates could be involved in mRNA maturation, with the DCD domain potentially recognizing m^5^C in mRNA, whereas the eTudor domain interacts with methylated Sm proteins, and might, in choanoflagellate species yet to be identified that encode an Argonaute protein(s) with N-terminal SDMA residues, interact with those as well. Consistent with this possibility, the m^5^C modification of mRNA appears to promote export from the nucleus (Yang et al. 2017), implicating eTudor-DCD proteins in trafficking of spliced, mature mRNA into the cytoplasm.

This putative function could be an evolutionary foundation for the piRNA pathway, given that the eTudor proteins involved in this pathway generally localize to perinuclear germ granules, which are associated with clusters of nuclear pore complexes (NPCs), and have been proposed to be extensions of the nuclear pore environment (Updike et al. 2011; Sengupta and Boag 2012; Sheth et al. 2010). These granules are foci of RNA and protein accumulation that appear to determine which mRNAs are permitted to enter the germ cell cytoplasm for translation, primarily, via silencing of unlicensed transcripts by Piwi-related Argonautes, and so are final arbiters of nuclear export (Yang et al. 2017; Voronina et al. 2011; Sheth et al. 2010; Sengupta and Boag 2012; Updike et al. 2011). Thus, choanoflagellate eTudor-DCD proteins, that can be predicted to interact with m^5^C and methylated Sm proteins during mRNA maturation and exit from the nucleus, could have been fundamental contributors to the function of the perinuclear granules that ultimately arose in animal germ cells and which also govern nuclear mRNA export by associating with NPCs (Sheth et al. 2010; Updike et al. 2011; Sengupta and Boag 2012). Given the role of piRNA pathway eTudor proteins in self vs. non-self discrimination, it is possible that eTudor-DCD proteins already play a role in this process in choanoflagellates, ensuring that self mRNAs are correctly spliced, modified, and licensed to exit the nucleus.

The ‘sudden’ appearance of a sizeable group of eTudor proteins and a fully-fledged piRNA pathway in the basal metazoans, the sponges, is now illuminated by the identification of a potential transitional form in choanoflagellates (Grimson et al. 2008; Fierro-Constaí;n et al. 2017). We propose that the choanoflagellate eTudor-DCD protein was an evolutionary bridge from Tudor-SN to the multi-eTudor proteins in metazoans that initiated the first piRNA pathway (Figure 14) (Jin et al. 2009; Caudy et al. 2003). The multi-eTudor proteins fused with a YTH domain we detected in several coral species provide additional support for this hypothesis because YTH and DCD are fused to each other in many species of chromist algae, the presumed donor of DCD to choanoflagellates. YTH domain proteins are known to localize to stress granules, which have considerable similarity to germ granules, where they concentrate mRNAs with the m^6^A modification to promote their translation (Voronina et al. 2011; Anders et al. 2018). YTH proteins additionally localize to nuclear speckles, also known as interchromatin granule clusters that are enriched in mRNA maturation factors, where they facilitate processing and nuclear export of m^6^A RNA (Galganski, Urbanek, and Krzyzosiak 2017). It is plausible that DCD domain proteins would similarly promote accumulation of modified mRNAs, and did so at the dawn of the piRNA pathway in the first metazoans. Such ancestral activity might underlie the germ granule localization of eTudor proteins and Piwi-related Argonautes complexed with piRNAs, which effectively concentrate RNA, via base pairing between mRNA and piRNA, as well as through binding of SDMA in the Argonaute

N-termini by multiple, fused eTudor domains. The activity of Piwi-related Argonaute and eTudor piRNA pathway factors has not been reported to involve mRNA modifications that might once have been recognized by DCD, or additional modifications by CAPAM-like methyltransferases, although that possibility merits further inquiry (Figure 14). The metazoan CAPAM-like methyltransferases could have originated from an ancestor present in a choanoflagellate eTudor-DCD protein, and mRNA modifications might still play a role in the piRNA pathway.

The predicted eTudor proteins from non-loricate choanoflagellates we analyzed are incomplete at their N-termini (Figure 13), and comparison with full length eTudor proteins from metazoans led us to speculate that the full length choanoflagellate proteins (that still reamin to be identified) contain N-terminal Znfs. Furthermore, we hypothesize that an evolutionary intermediate eTudor protein existed in the first metazoans which contained multiple eTudor domains and an N-terminal Znf, but lacked the DCD domain, its function replaced by interaction with Piwi-related Argonautes and their associated small RNAs (Figure 14). Although the variety of the Znfs in this founding member is difficult to determine with the available data, a protein with this basic architecture could have served as a foundation for the evolution of the piRNA pathway, ultimately, derived from an ancestral choanoflagellate eTudor-DCD protein. Our survey of metazoan eTudor proteins also revealed a divergent evolutionary trajectory in nematodes which lack eTudor proteins with N-terminal Znfs, or with more than two copies of eTudor, a apparently occurred. A fundamental contributing factor to this outcome was a deletion of the conserved α-helix in one of the eTudor domains in the ancestor of the tandem eTudor proteins that are part of the nematode-specific RNA-dependent RNA polymerase (RdRP) complexes (Supplementary Figure 10).

### Conclusions

Our comprehensive search for EVE domains demonstrated their wide presence in bacteria and archaea, as well as most eukaryotes. Mechanistically, the common denominator of the EVE family is the binding of methylated bases in DNA and RNA that turns out to be important in a broad variety of functional contexts. The (predicted) biological roles of EVE-like domains are many, and depending on the context, might seem to be in opposition. However, the overarching theme is involvement in immunity, self vs non-self discrimination, and stress response/PCD, whether that be through targeting of modified DNA for restriction in diverse prokaryotes, regulation of the proliferation-PCD-dormancy balance in Proteobacteria and eukaryotes (especially during the differentiation of immune cell populations in vertebrates), or export and translation of spliced and modified mRNAs in choanoflagellates.

The linkage between EVE proteins, tRNA modification and cytochrome c maturation that is suggested by the conserved operonic organization in α-proteobacteria could shed light on the mechanism of the reported anti-apoptotic action of eukaryotic THYN1/Thy28-like EVE proteins and, beyond that, the origin of cytochrome c involvement in eukaryotic PCD. We hypothesize that the proteins encoded in the α-proteobacterial TsaD->GpsA->YciI->EVE operon promote translation coupled with respiration, and down-regulation of this operon could induce dormancy. It is unclear if cytochrome c efflux occurs in free-living α-proteobacteria undergoing dormancy or PCD as it does in mitochondria, or this phenomenon evolved during eukaryogenesis, but it would likely be an effective mechanism for rapidly decreasing ATP production in the event of a runaway viral infection.

The EVE domains split into two functionally distinct classes that evolve under different regimes, slowly in the case of those that seem to be involved in basic cellular functions in Proteobacteria and eukaryotes, and fast, in the case of those involved in defense functions and virus-host arms races in diverse bacteria and archaea. The incorporation of EVE domains into numerous RM and TA modules is a remarkable, previously unnoticed pattern in microbial evolution, which emphasizes the various, still under-appreciated roles that different base modifications play in the intricate virus-host interactions.

The DCD domains present another facet of the PUA/EVE story. The genomic context in prokaryotes implies that they perform functions similar to those of the second class of EVE domains, namely, are involved in antivirus defense via recognition of modified bases in DNA. However, in eukaryotes, the DCD domains become part of a more complicated evolutionary scenario that seems to involve an important aspect of the origins of plants and animals. Plants have retained the DCD domain in multiple proteins that contribute to PCD during plant development as well as stress and pathogen response. In animals, DCD domains apparently have been lost. However, we identified a ‘smoking gun’ in choanoflagellates, the unicellular direct ancestors of animals, where the DCD domains are fused to eTudor proteins. The roles played by eTudor proteins in germline immortality, gametogenesis, and early embryonic development in animals, where regulation of PCD is pivotal (Schrader, Kaldenhoff, and Richter 1997), are intriguingly similar to those of DCD proteins in plants. It appears likely, therefore, that DCD domains were important at the earliest stages of the evolution of multicellularity in both plant and animal ancestors, but then, were supplanted by the expanding eTudor family in the animal lineage.

The extent to which the regulation of PCD might still be relevant to eTudor and piRNA function, despite the loss of the DCD domain, remains to be elucidated, but there is evidence suggesting that it could be considerable. In the light of the putative evolutionary connection between the ancient form of PCD and the evolution of the eTudors and the piRNA network, it seems more explicable that conserved piRNA biogenesis factors Tudor-KH and MitoPLD/Zucchini are mitochondrially localized, perinuclear piRNA processing germ granules are also closely associated with mitochondria in addition to nuclear pores, and that reported phenotypes of many piRNA factor mutants include induction of PCD and/or germline mortality (Bronkhorst and Ketting 2018; Weick and Miska 2014; Houwing et al. 2007; Tóth et al. 2016; Aravin and Chan 2011; Manage et al. 2020). PCD is an important part of tissue differentiation, and piRNAs can prevent ectopic expression of somatic genes that might contribute to PCD or senescence of the germline (Strome and Updike 2015; Meier, Finch, and Evan 2000). In animals, populations of piRNAs, with their biogenesis orchestrated by eTudor proteins, Piwi-related Argonautes and other factors, could have taken the place of the DCD domain in regulating PCD and transposable element mobilization by licensing germline mRNA translation. Whereas in ancestral eukaryotes, m^5^C modifications of RNA possibly regulated nuclear export in the absence of a piRNA pathway, when they began to evolve a germline and differentiation into tissues from an embryo in the metazoan lineage, Argonaute-piRNA complexes may have taken over aspects of this screening process, ultimately linking it to Argonaute-directed repressive chromatin modification, a conserved feature of eukaryotic cells, in order to regulate transcription as well as trafficking from the nucleus (Matzke and Mosher 2014; Wälti et al. 2006; Law and Jacobsen 2010). The consistent presence of ubiquitin-associated Znf domains in eTudor factors suggests an additional role in protein degradation, interconnecting multiple layers of control over expression.

The DCD and YTH domains have strikingly similar evolutionary histories and functional associations. Both are found only rarely in prokaryotes, mostly in archaea, where they are associated with restriction of modified DNA. PSI-BLAST searches for both DCD and YTH domains in prokaryotes readily recover restriction-associated EVE domain proteins, implying that they are both essentially varieties of the much more numerous EVE domains, produced from the diversification driven by virus-host conflict. Subsequently, it would appear, during eukaryogenesis, both were plucked from relative obscurity and conscripted into conserved roles in eukaryotes where they are (possibly, in the case of DCD) involved with concentration and processing of modified RNA, especially during stress response/PCD. The piRNA system, born of the partnership of eTudor proteins and Piwi-related Argonautes in the first animals and showing parallels with DCD function in plants, is conceivably related to this ancient RNA concentration and processing mechanism, and possibly, still involves RNA modifications. Consequently, characterization of the roles of choanoflagellate eTudor-DCD proteins as well as metazoan eTudor proteins in PCD and stress response, could be important for understanding the origins of animal germline specification. Moreover, the study of restriction systems with EVE-like domains is likely to shed light on the origins of eukaryotic mRNA regulation.

Taken together, the findings reported here suggest multiple connections between PCD, antivirus defense, and various forms of stress response via diverse families of EVE-like domains that recognize modified bases in DNA and RNA. The role of such bases in the coordination of antivirus defense, PCD, cell proliferation and development remains under-appreciated, perhaps, substantially. These observations open up many experimental directions that can be expected to advance the understanding of the complexity of all these processes.

## Materials and Methods

### Identification and phylogenetic analysis of EVE and DCD domain proteins

A comprehensive search for EVE proteins was initiated with the available multiple sequence alignment pfam01878. This alignment was clustered and each sub-alignment used to produce a position-specific scoring matrix (PSSM) for use as a PSI-BLAST query against the non-redundant (nr) NCBI database (E value ≤ 10) (Altschul et al. 1997). Manual filtering of these results aided by BLAST and HHpred (Zimmermann et al. 2018) was followed by extraction of the EVE domains from each protein and similarity clustering to a threshold of 0.85 with MMeqs2 (Hauser, Steinegger, and Söding 2016).

Selection of one representative per cluster yielded 8,403 sequences for CLANS analysis (Frickey and Lupas 2004) and phylogenetic tree construction. The metrics of within-group similarity in Proteobacteria were calculated using a tree built with the FastTree program (Price, Dehal, and Arkin 2010) from an alignment of these sequences made using PROMALS3D (Pei and Grishin 2007). This tree was rooted at the midpoint, after which all root-to-tips distances were calculated, giving a median tree height of 2.82 with and interquartile range of 2.27-4.28. The subtrees for α-proteobacteria and β/γ-proteobacteria were extracted and the same values were calculated, yielding heights of 0.99 [0.85-1.18] and 0.9 [0.66-1.03], respectively, which represent 35% and 32% of the full tree median height. During the generation of the schematic tree included in the Supplementary Data, the sequences were further clustered to a similarity threshold of 0.5 with MMseqs2 (Hauser, Steinegger, and Söding 2016), then the sequences in each cluster were aligned with MUSCLE (Edgar 2004). Next, profile-to-profile similarity scores between all clusters were calculated with HHsearch (Söding 2004), and a UPGMA dendrogram was generated using the pairwise similarity scores. Clusters with high similarity, defined as a pairwise score to self-score ratio > 0.1, were aligned to each other with HHalign (Söding et al. 2006). This procedure was performed for a total of 5 iterations. Finally, each cluster alignment was used to build trees using the FastTree program (Price, Dehal, and Arkin 2010) that were rooted at the midpoint and grafted onto the tips of the UPGMA dendrogram that was generated from the cluster similarity scores.

Searches for DCD and eTudor domains were performed similarly to the EVE domains, using the pfam10539 and pfam00567 alignments. For the choanoflagellate eTudor-DCD proteins, a BLAST database was constructed using the predicted protein sequences published by Richter et al. (Richter et al. 2018) prior to searching with PSI-BLAST. Multiple sequence alignments of EVE, DCD, and eTudor domains were constructed with MUSCLE and PROMALS3D (Pei and Grishin 2007; Edgar 2004).

### Genome neighborhood analysis

Domains in genes neighboring prokaryotic EVE and DCD genes (10 on each side) were identified using PSI-BLAST against alignments of domains in the NCBI Conserved Domain Database (CDD) and Pfam (E-value 0.001). Some genes were additionally analyzed manually using HHpred. Gene sequence, coordinate and directional information was downloaded from GenBank using custom Perl scripts. The contextual information network graph was generated using the Cytoscape program (Shannon et al. 2003). The thickness of edges between nodes represents the strength of association between domains. For any pair of domains in a given genomic neighborhood, the association was calculated as 1/(n+1) where n is the number of intervening domains encoded in the neighborhood by distinct genes between the members of the pair. The association values were first averaged across the neighborhoods within each genus and then averaged between the genera to produce the overall weighted average.

## Acknowledgements

The authors thank Dr. Andrew Z. Fire (Stanford University) for critical reading of the manuscript and insightful comments, and Koonin group members for helpful discussions.

## Declarations

### Availability of data and materials

The datasets supporting the analysis presented in this article are included within the additional files.

### Authors’ contributions

Conception and design of the study: RB, EVK. Data collection: RB. Data analysis: RB, YIW, EVK. Manuscript drafting: RB, EVK. Manuscript revision for critical intellectual content: RB, EVK. All authors read and approved the final manuscript.

### Consent for publication

Not applicable.

### Ethics approval and consent to participate

Not applicable.

