## Supplementary figures for "Modified base-binding EVE and DCD Domains Implicated in the Origins of Programmed Cell Death and the piRNA Pathway"

**Table of Contents**


| Page | Figure title |
|---|---|
| 1 | **Supplementary Figure 1: Phylogenetic distribution of prokaryotic EVE proteins** |
| 2 | **Supplementary Figure 2: Conserved genomic context of EVE proteins in Vibrionales** |
| 2 | **Supplementary Figure 3: Conserved genomic context of EVE proteins in Alteromonadales** |
| 3 | **Supplementary Figure 4: Conserved genomic context of EVE proteins in Pasteurellales** |
| 4 | **Supplementary Figure 5: Schematic phylogenetic tree of EVE domain cluster representatives** |
| 5 | **Supplementary Figure 6: Conserved genomic context of EVE proteins in *Nocardia* and related genera that are found in a distinct CLANS analysis cluster (green in Figure 2)** |
| 5 | **Supplementary Figure 7: Conserved genomic context of EVE proteins in *Azospirillum* that are found in a distinct CLANS analysis cluster (green in Figure 2)** |
| 6 | **Supplementary Figure 8: COG1743->DUF499->SWI2/SNF2 helicase-nuclease->EVE defense systems in archaea** |
| 7 | **Supplementary Figure 9: Alignment of EVE and DCD domain representatives** |
| 8 | **Supplementary Figure 10: Hypothesis for the evolution of the RdRP complex eTudor proteins in *C. elegans*** |
| 9 | **Supplementary Figure 11: Alignment of the eTudor domain from *Drosophila* SND1 (Tudor-SN) with eTudor domains in choanoflagellates that are fused to DCD domains** |
| 10 | **Supplementary Reference** |

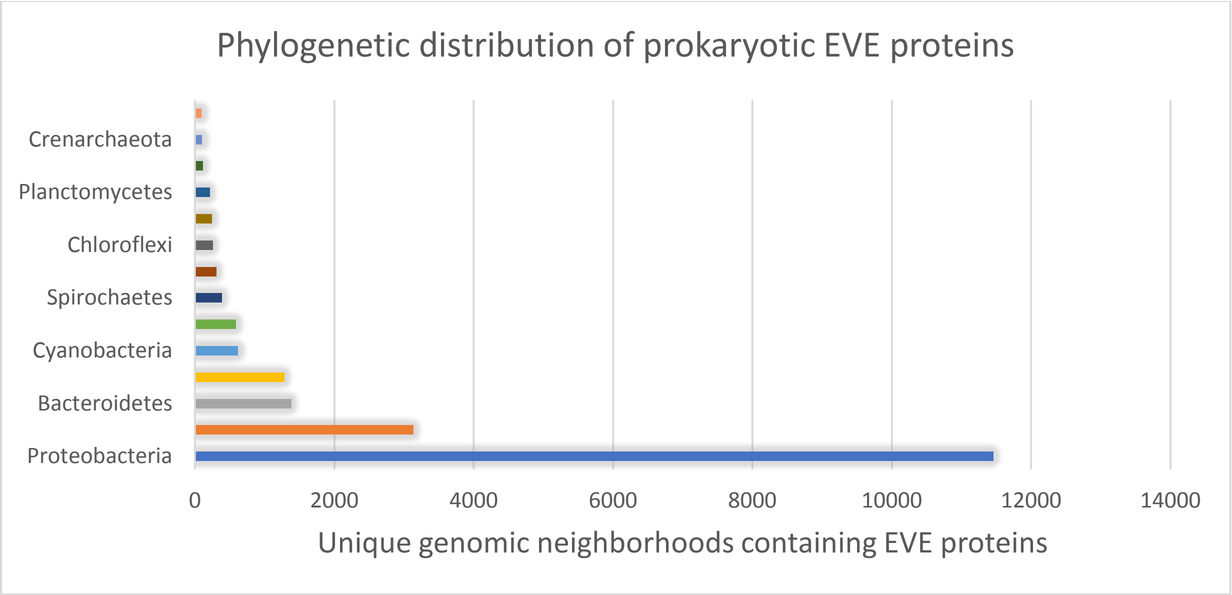


**Supplementary Figure 1, Phylogenetic distribution of prokaryotic EVE proteins**

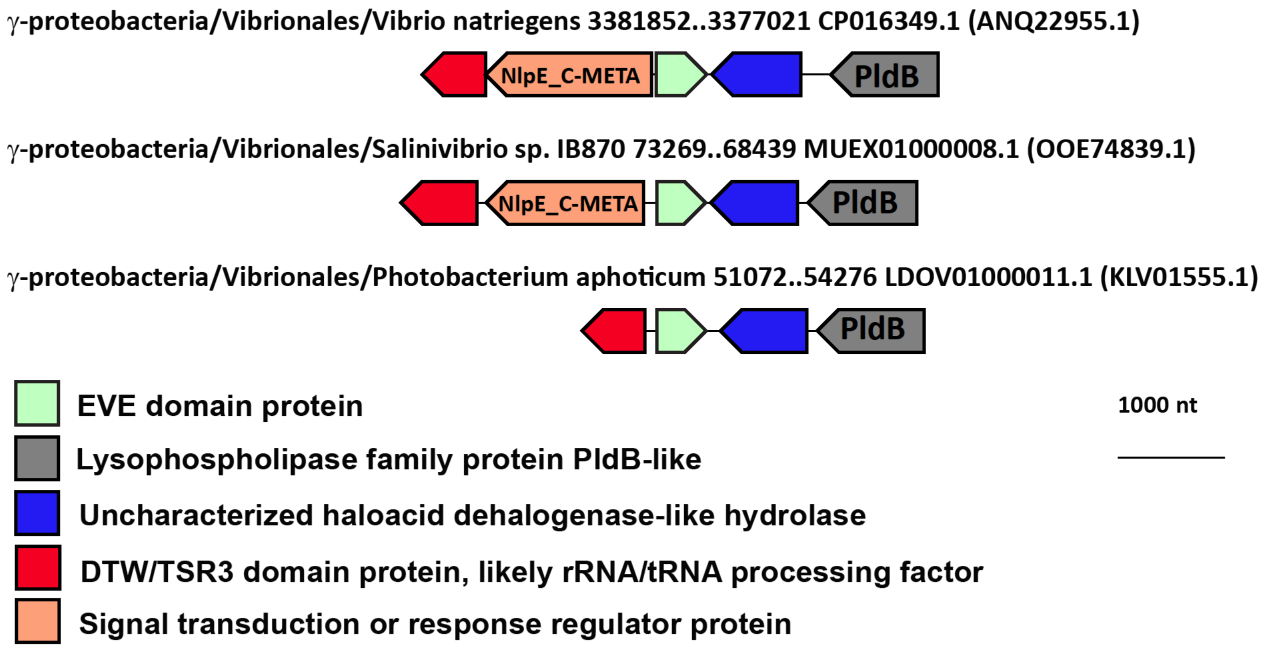


#### Supplementary Figure 2, Conserved genomic context of EVE proteins in Vibrionales

Representative EVE protein neighborhoods from Vibronales. Genes are shown as arrows from 5’ to 3’. The order of α-proteobacteria, species, and genomic coordinates for each neighborhood are indicated, as are the GenBank genome accessions and, in parentheses, the GenBank accessions for each EVE protein.

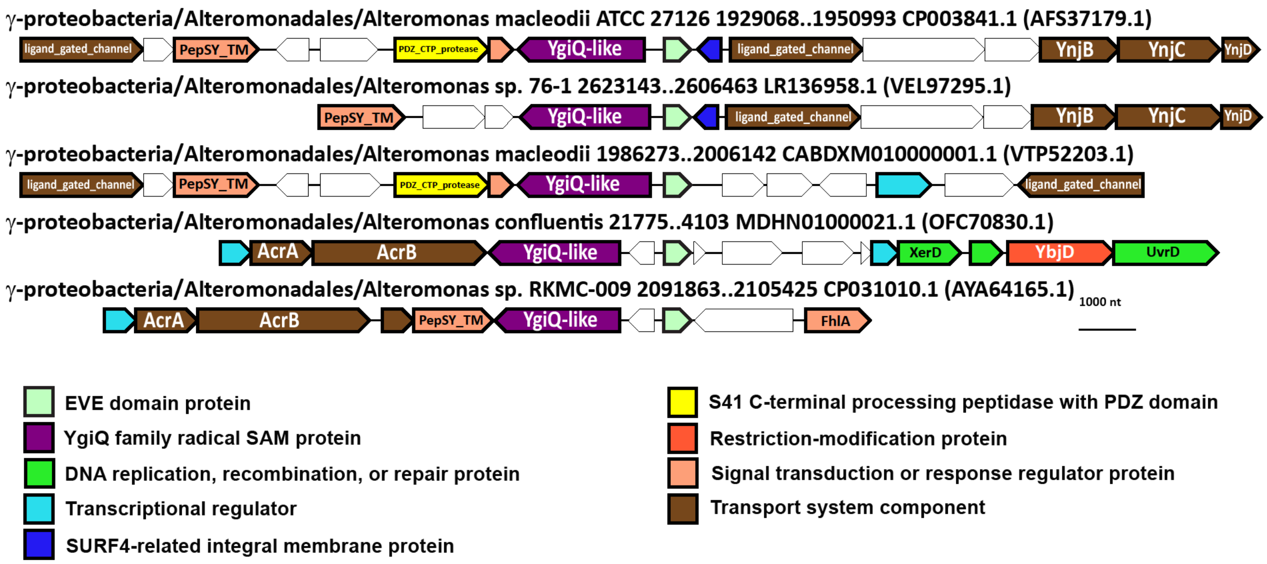


#### Supplementary Figure 3, Conserved genomic context of EVE proteins in Alteromonadales

Representative EVE protein neighborhoods from Alteromonadales. Genes are shown as arrows from 5’ to 3’. The order of α-proteobacteria, species, and genomic coordinates for each neighborhood are indicated, as are the GenBank genome accessions and, in parentheses, the GenBank accessions for each EVE protein.

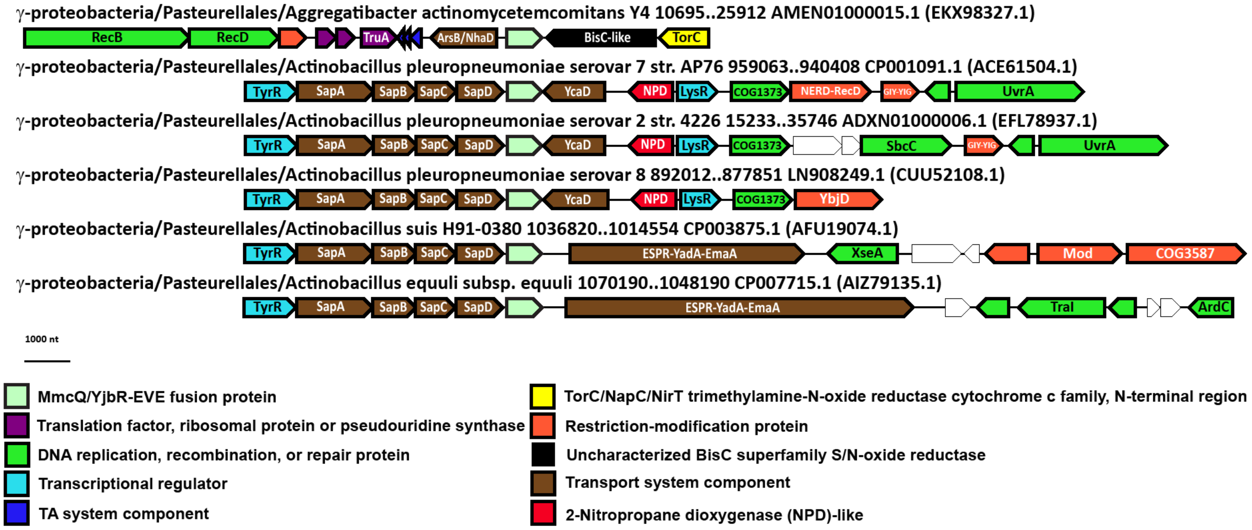


### Supplementary Figure 4, Conserved genomic context of MmcQ/YjbR-EVE fusion proteins in Pasteurellales

Representative MmcQ/YjbR-EVE fusion protein neighborhoods from Pasteurellales. Genes are shown as arrows from 5' to 3'. The order of α-proteobacteria, species, and genomic coordinates for each neighborhood are indicated, as are the GenBank genome accessions and, in parentheses, the GenBank accessions for each EVE protein.

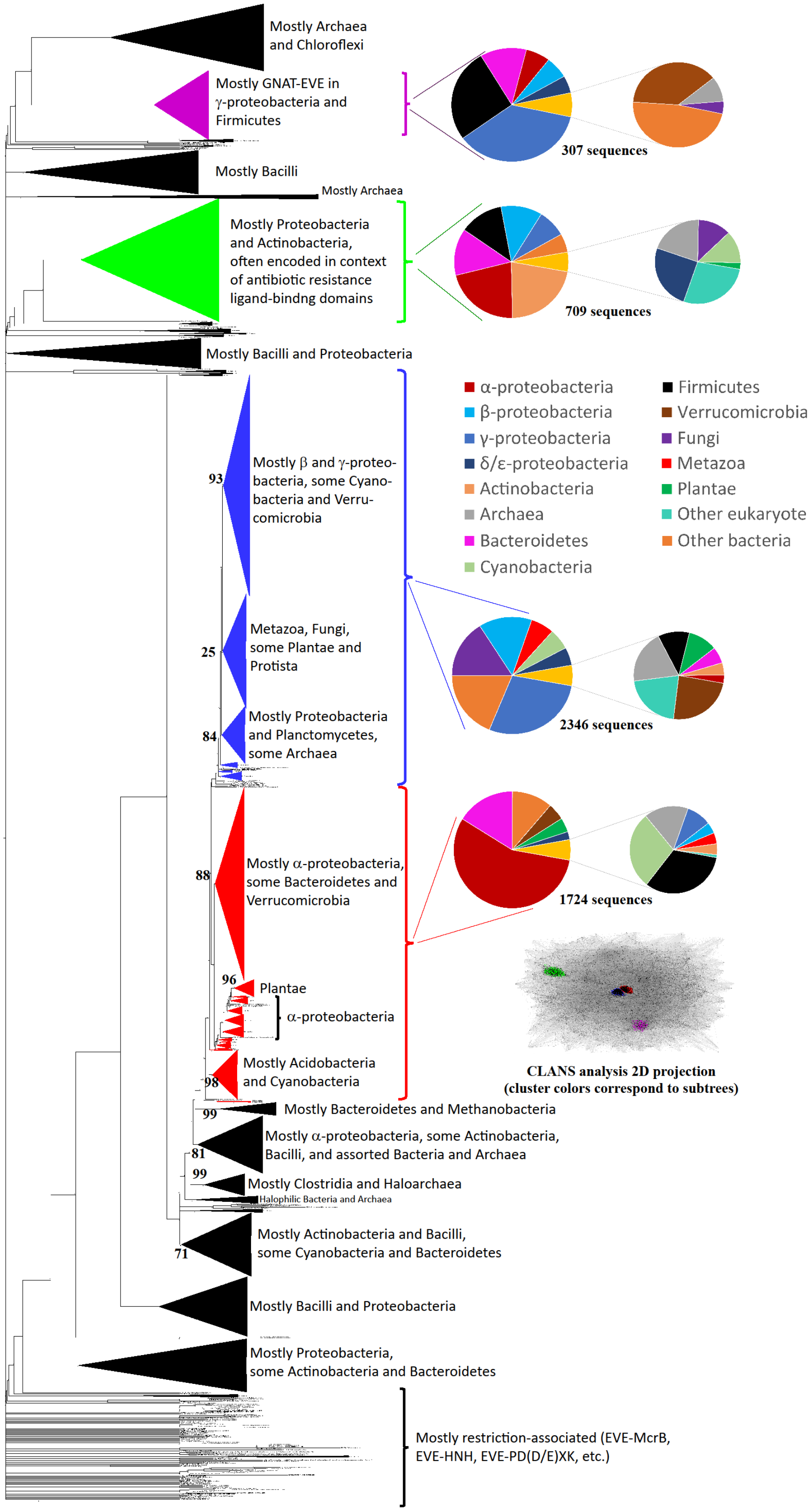


**Supplementary Figure 5, Schematic phylogenetic tree of EVE domain cluster representatives**

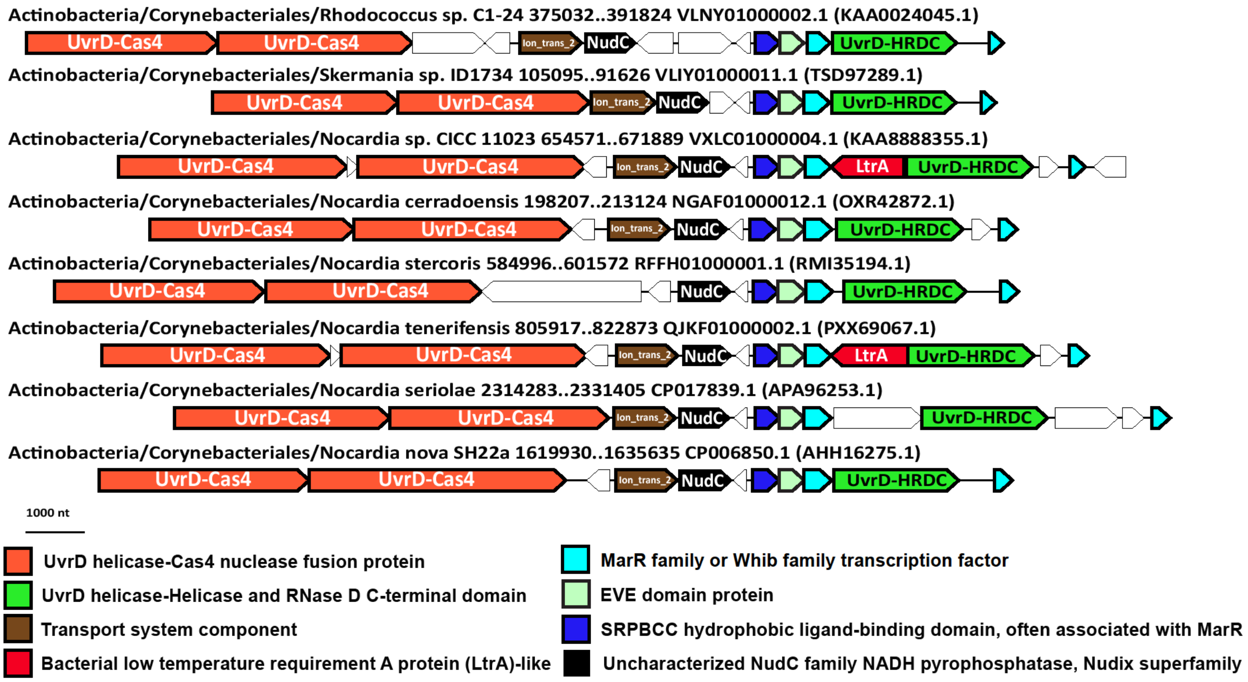


### Supplementary Figure 6, Conserved genomic context of EVE proteins in *Nocardia* and related genera that are found in a distinct CLANS analysis cluster (green in Figure 2)

Representative EVE protein neighborhoods from *Nocardia* and related genera. Genes are shown as arrows from 5' to 3'. The order of α-proteobacteria, species, and genomic coordinates for each neighborhood are indicated, as are the GenBank genome accessions and, in parentheses, the GenBank accessions for each EVE protein.

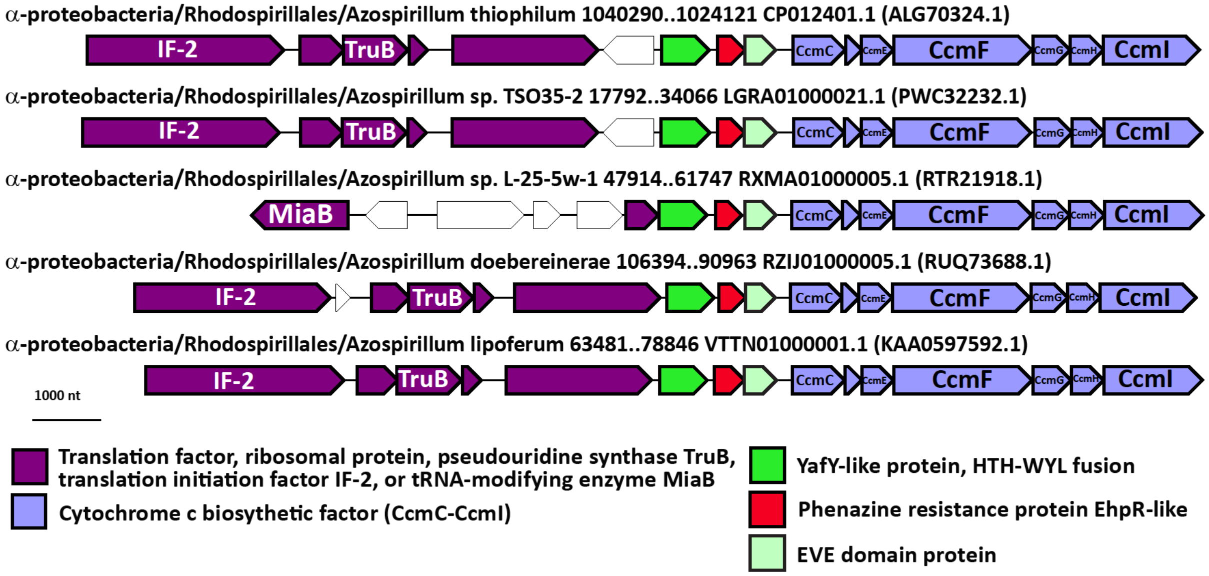


### Supplementary Figure 7, Conserved genomic context of EVE proteins in *Azospirillum* that are found in a distinct CLANS analysis cluster (green in Figure 2)

Representative EVE protein neighborhoods from *Azospirillum*. Genes are shown as arrows from 5' to 3'. The order of α-proteobacteria, species, and genomic coordinates for each neighborhood are indicated, as are the GenBank genome accessions and, in parentheses, the GenBank accessions for each EVE protein.

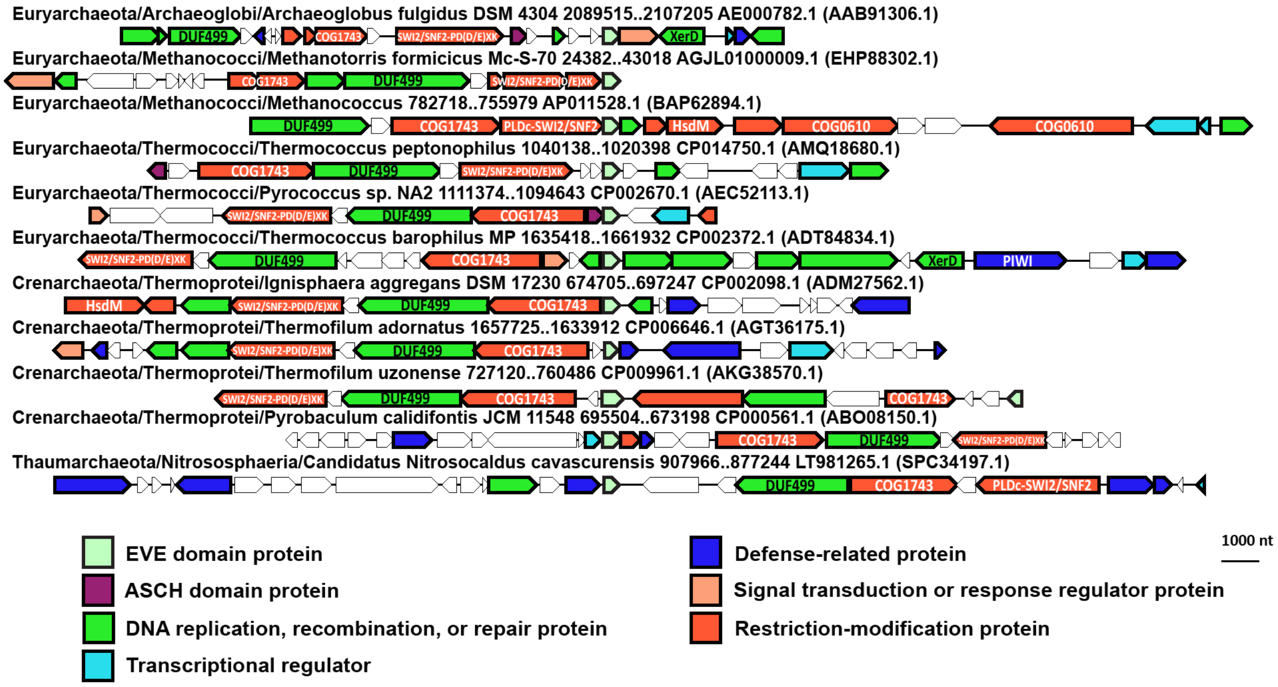


**Supplementary Figure 8, COG1743->DUF499->SWI2/SNF2 helicase-nuclease->EVE defense systems in diverse archaea**

EVE protein neighborhoods from archaea that contain COG1743->DUF499->SWI2/SNF2 helicase-nuclease operons. Genes are shown as arrows from 5' to 3'. The order of α-proteobacteria, species, and genomic coordinates for each neighborhood are indicated, as are the GenBank genome accessions and, in parentheses, the GenBank accessions for each EVE protein.

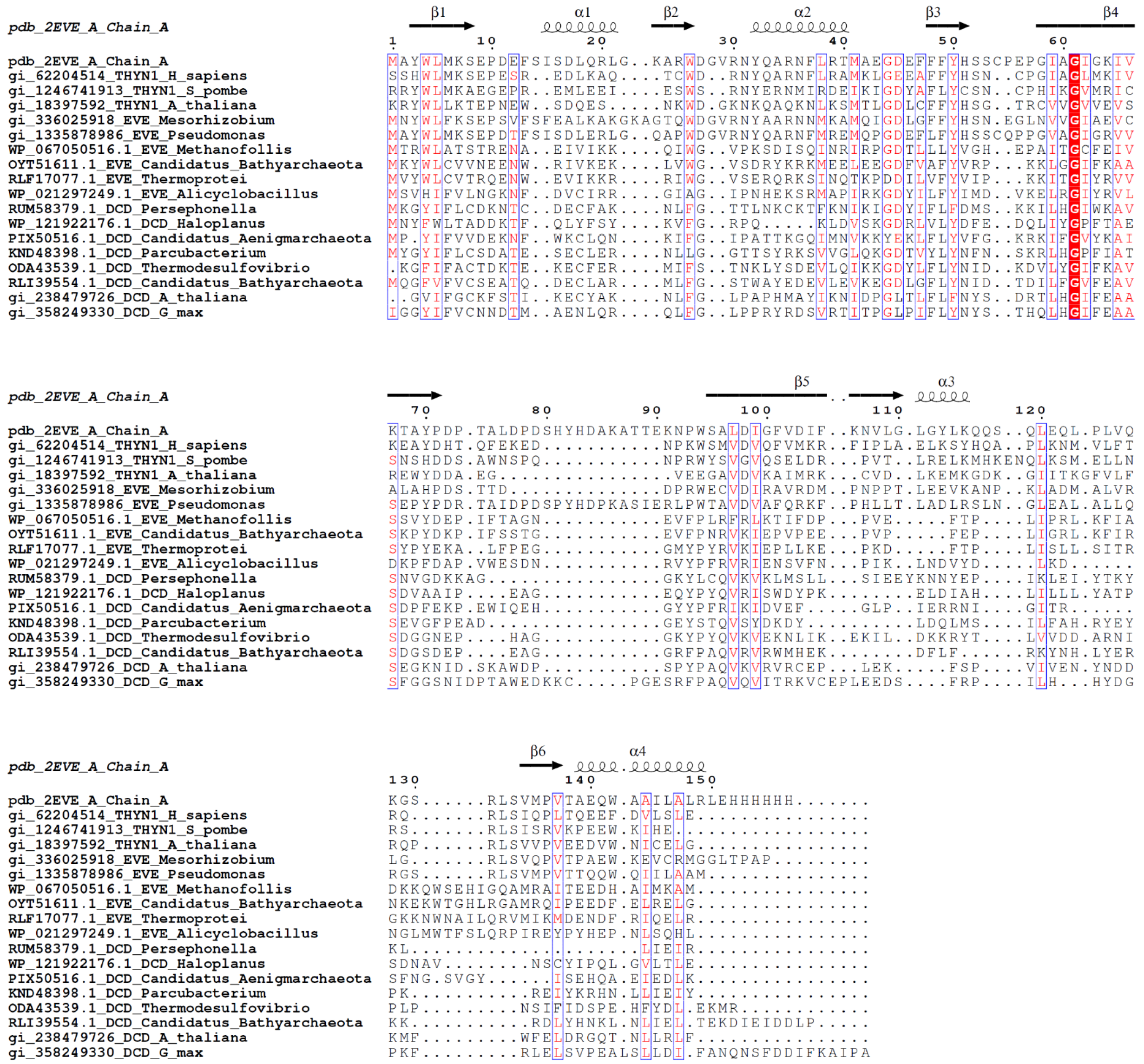


### Supplementary Figure 9, Alignment of EVE and DCD domains

Alignment of representative EVE and DCD domain sequences. The alignment was made with PROMALS3D and rendered with ESPript (Robert and Gouet 2014). The first 6 sequences are highly conserved EVE domains of the type found in the two largest CLANS clusters, followed by 3 which are more similar to DCD domains. The final 8 sequences are DCD domains, and the last two are canonical DCD domains from plants, including the most characterized from N-rich protein (NRP) in *Glycine max*.

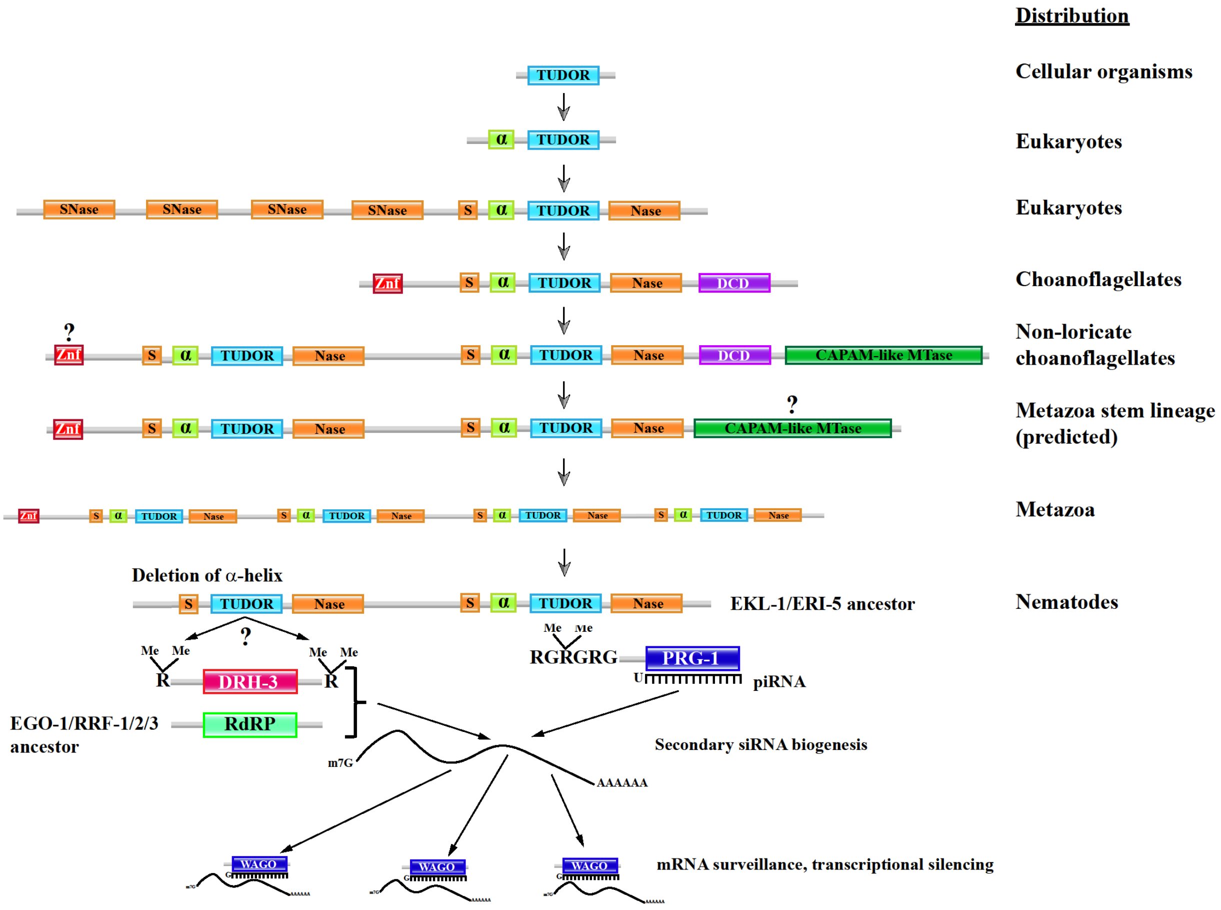


**Supplementary Figure 10, Hypothesis for the evolution of the RdRP complex eTudor proteins in *C. elegans***

In *C. elegans*, the piRNA pathway has become hyperdeveloped and integrated with the Dicer-dependent RNAi response. The foundation for this is apparently the mutation of an eTudor domain in a tandem eTudor protein that was the ancestor of the EKL-1 and ERI-5 eTudor proteins. These eTudor factors are core components of distinct RNA dependent RNA polymerase (RdRP) complexes responsible for secondary siRNA biogenesis in worms, which occurs in response to binding of mRNA by Dicer-dependent primary siRNAs generated from dsRNA as well as piRNAs, which in *C. elegans* are transcribed from individual loci. The other factors in the complexes minimally include an RdRP and DRH-3 (Dicer-related helicase-3). Presumably, the mutation in the N-terminal eTudor domain allows for a novel type of interaction with a methylated protein, likely one of the other RdRP complex components. We predict this factor to be DRH-3, as it is present in all RdRP complexes, although it could be the various RdRPs that distinguish different RdRP complexes, or another factor altogether.

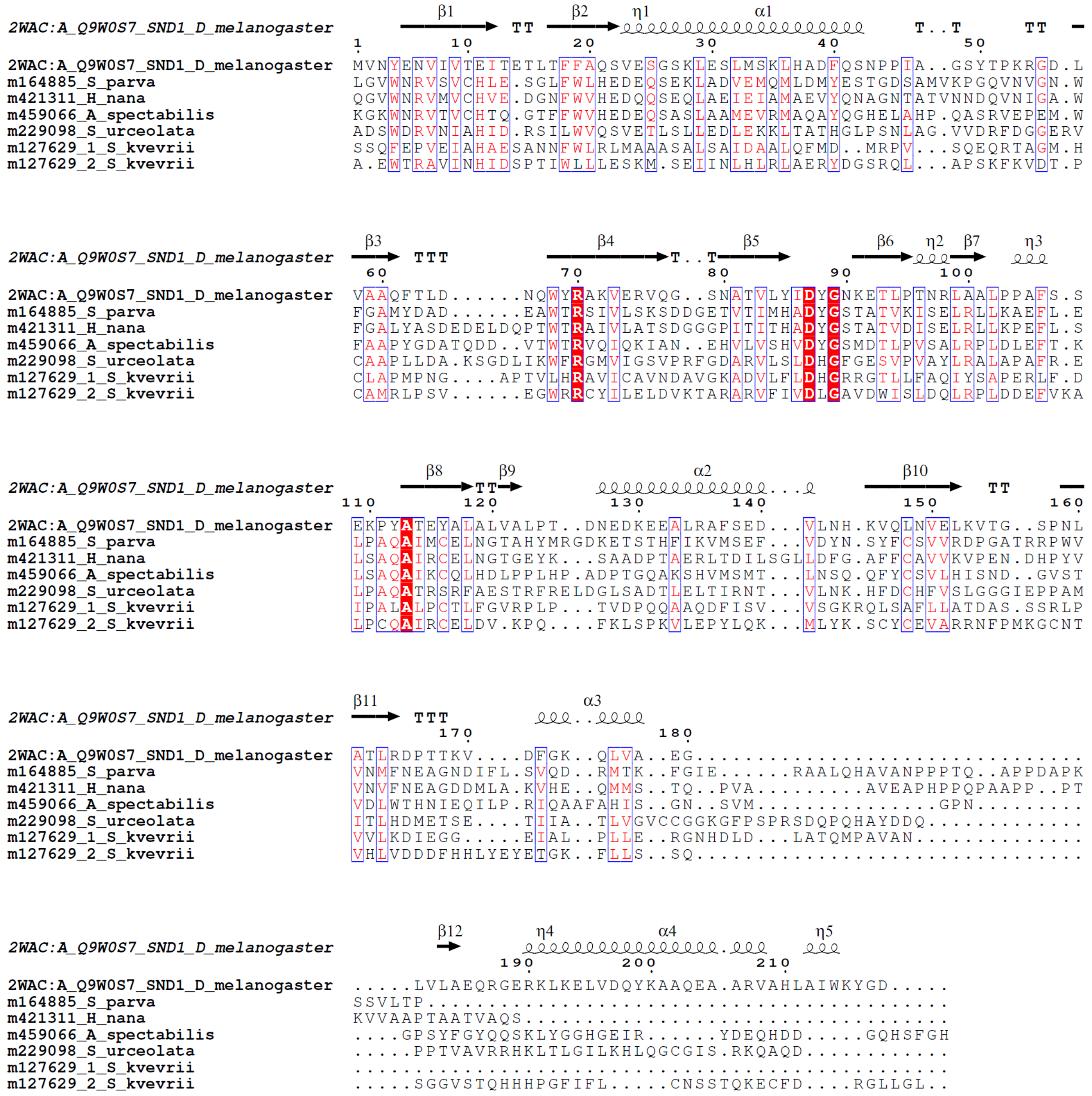


### Supplementary Figure 11, Alignment of eTudor domains in choanoflagellates that are fused to DCD domains and the eTudor domain from *Drosophila* SND1 (Tudor-SN)

Alignment of eTudor domain sequences from choanoflagellates that are found in DCD proteins and the eTudor domain from the Tudor-SN ortholog encoded by *D. melanogaster*, SND1. The alignment was made with PROMALS3D and rendered with ESPript (Robert and Gouet 2014). The conserved residues and secondary structures characteristic of eTudor domains are present.
